# Tumor Necrosis Factor induces pathogenic programmed macrophage necrosis in tuberculosis through a mitochondrial-lysosomal-endoplasmic reticulum circuit

**DOI:** 10.1101/511436

**Authors:** Francisco J. Roca, Sarah Redmond, Lalita Ramakrishnan

## Abstract

Necrosis of infected macrophages constitutes a critical pathogenetic event in tuberculosis releasing mycobacteria into the extracellular environment where they can grow unrestricted. In zebrafish infected with *Mycobacterium marinum*, a close relative of *Mycobacterium tuberculosis*, excess Tumor Necrosis Factor triggers programmed necrosis of infected macrophages through the production of mitochondrial reactive oxygen species (ROS) and the participation of cyclophilin D, a component of the mitochondrial permeability transition pore. Here we show that this necrosis pathway is not mitochondrion-intrinsic but rather results from an interorganellar circuit initiating and culminating in the mitochondrion. Mitochondrial ROS induce production of lysosomal ceramide which ultimately activates the cytosolic protein BAX. BAX promotes calcium flow from the endoplasmic reticulum into the mitochondrion through ryanodine receptors. The resultant mitochondrial calcium overload triggers cyclophilin D-mediated necrosis. We identify ryanodine receptors and plasma membrane L-Type calcium channels as specific druggable targets to intercept mitochondrial calcium overload so as to inhibit macrophage necrosis.

## INTRODUCTION

The pathogenic life cycle of *Mycobacterium tuberculosis* (Mtb), the agent of human tuberculosis is fueled by its multiple interactions with host macrophages (Cambier et al., 2014; Ramakrishnan, 2012; Srivastava et al., 2014). The bacteria use macrophages to traverse host epithelial barriers to enter deeper tissues, whence they recruit additional macrophages to form granulomas, pathognomonic structures that can serve as intracellular bacterial growth niches (Cambier et al., 2014; Cohen et al., 2018; Ramakrishnan, 2012; Srivastava et al., 2014). Granuloma macrophages can undergo necrosis, a key pathogenic event that further increases bacterial growth in the more permissive extracellular milieu (Behar et al., 2010; Chen et al., 2008; Divangahi et al., 2013; Divangahi et al., 2009; Ramakrishnan, 2012), thereby increasing both disease morbidity and transmission (Bekker and Wood, 2010; Cambier et al., 2014; Huang et al., 2014; Reichler et al., 2002).

Mycobacterium-macrophage interactions and resultant macrophage fates can be detailed in the optically transparent zebrafish larva infected with *Mycobacterium marinum* (Mm), a close genetic relative of Mtb (Pagan and Ramakrishnan, 2014; Takaki et al., 2013). In this model, distinct host genetic mutations that increase macrophage necrosis all render the host hypersusceptible by promoting unrestricted extracellular mycobacterial growth (Berg et al., 2016; Clay et al., 2008; Pagan et al., 2015; Tobin et al., 2012). A notable genetic perturbation that produces hypersusceptibility through macrophage necrosis is the increased expression of leukotriene A4 hydrolase (LTA4H) that catalyzes the final step in the synthesis of the inflammatory lipid mediator leukotriene B4 (LTB_4_) (Tobin et al., 2012). Humans with a common functional LTA4H promoter variant that increases its expression are also hypersusceptible to TB (Thuong et al., 2017; Tobin et al., 2012). Among a case cohort of tuberculous meningitis, the severest form of TB, LTA4H-high individuals had increased risk of death. Consistent with excessive inflammation being the cause of death, survival was dramatically increased among patients who received adjunctive anti-inflammatory therapy with glucocorticoids (Thuong et al., 2017; Tobin et al., 2012).

The human relevance of the zebrafish findings provided the impetus to carry out a detailed mechanistic dissection of the necrosis pathway. Returning to the zebrafish, we showed that susceptibility of the LTA4H-high state is due to the excessive production of the pro-inflammatory cytokine Tumor Necrosis Factor (TNF) (Tobin et al., 2012). We found that excess TNF triggers RIPK3-dependent programmed necrosis of infected macrophages (Galluzzi et al., 2012; Roca and Ramakrishnan, 2013). TNF-RIPK3 interactions increase mitochondrial reactive oxygen species (ROS) production which are required for necrosis along with cyclophilin D, a mitochondrial matrix protein (Roca and Ramakrishnan, 2013). Oxidative stress is a reported trigger for the activation of cyclophilin D, which causes necrosis by promoting sustained opening of the mitochondrial permeability transition pore complex (MPTP) thereby leading to mitochondrial swelling, loss of membrane potential, disruption of the outer membrane and ATP depletion, (Baines et al., 2005; Bernardi, 2013; Bernardi et al., 2006; Du and Yan, 2010; Halestrap, 2005; Halestrap et al., 2004; Halestrap et al., 1998; Nakagawa et al., 2005). Accordingly, the dual requirement for mitochondrial ROS and cyclophilin D was consistent with a mitochondrion-intrinsic pathway where TNF-induced mitochondrial ROS activate cyclophilin D (Figure 1A).

**Figure 1:**
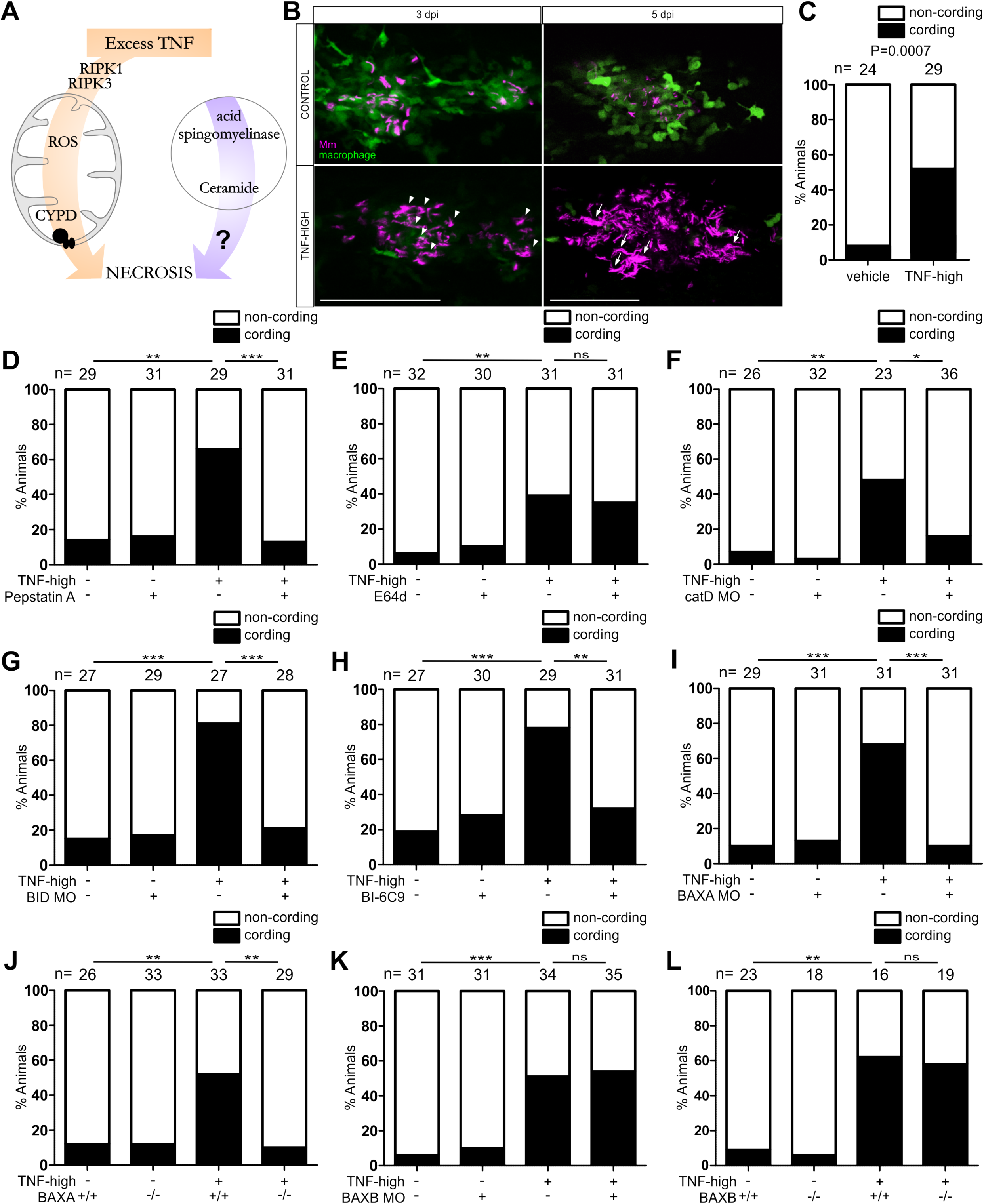
Ceramide contributes to cell death by engaging catD, BID and BAX. (A) Cartoon showing previously identified components of the pathway by which excess TNF triggers programmed necrosis in mycobacterium-infected macrophages. CYPD, cyclophilin D. (B) Confocal microscopy of granulomas in 3 dpi or 5 dpi *Tg(mpeg1:YFP)*^*w200*^ larvae injected with TNF or vehicle. White arrowheads show extracellular bacteria. White arrows show extracellular bacteria with cording phenotype. A single z image is showed for the control image 5 dpi to appreciate cellularity and maximum intensity projection for TNF-high to appreciate the cording phenotype. Scale bar 100 μm. (C) Percentage of animals with cording among sibling larvae injected with TNF or vehicle. (Fisher’s exact test). Representative of 6 independent experiments. (D) Percentage of animals with cording among sibling larvae injected with TNF or vehicle in presence or absence of 5 μM Pepstatin A. **p < 0.01; ***p < 0.001 (Fisher’s exact test). Representative of 3 independent experiments. (E) Percentage of animals with cording among sibling larvae injected with TNF or vehicle in presence or absence of 1 μg/ml E64d. **p < 0.01 (Fisher’s exact test). Representative of 3 independent experiments. (F) Percentage of animals with cording among WT and cathepsin D morphant siblings injected with TNF or vehicle. *p < 0.05; **p < 0.01 (Fisher’s exact test). Representative of 4 independent experiments. (G) Percentage of animals with cording among WT and BID morphant siblings injected with TNF or vehicle. ***p < 0.001 (Fisher’s exact test). Representative of 4 independent experiments. (H) Percentage of animals with cording among sibling larvae injected with TNF or vehicle in presence or absence of 5 μM BI-6C9. **p < 0.01; ***p < 0.001 (Fisher’s exact test). Representative of 6 independent experiments. (I) Percentage of animals with cording among WT and BAXA morphants siblings injected with TNF or vehicle. ***p < 0.001 (Fisher’s exact test). Representative of 3 independent experiments. (J) Percentage of animals with cording among WT and BAXA mutant larvae injected with TNF or vehicle. **p < 0.01 (Fisher’s exact test). Representative of 6 independent experiments. (K) Percentage of animals with cording among WT and BAXB morphants siblings injected with TNF or vehicle. ***p < 0.001 (Fisher’s exact test). Representative of 2 independent experiments. (L) Percentage of animals with cording among WT and BAXB mutant larvae injected with TNF or vehicle. **p < 0.01 (Fisher’s exact test). Representative of 3 independent experiments.

However, necrosis also required lysosomal components, specifically ceramide produced by lysosomal acid sphingomyelinase (ASMase) (Roca and Ramakrishnan, 2013) (Figure 1A). In this work, we sought to understand why lysosomal ceramide would be required to complete mitochondrially mediated necrosis (Figure 1A). We show that TNF-mediated necrosis results from a single pathway, an interorganellar circuit that initiates in the mitochondrion and traverses the lysosome, cytosol, and endoplasmic reticulum (ER) before returning to the mitochondrion to finally execute necrosis. This intricate circuit that begins with mitochondrial ROS production ultimately results in mitochondrial Ca2+ overload, which is the likely trigger for cyclophilin D activation (Clarke et al., 2002; Halestrap, 2005; Halestrap et al., 2004; Halestrap et al., 1998; Orrenius et al., 2003). Elucidating this pathway has led to the identification of multiple commonly-used drugs that can inhibit TNF-induced necrosis by preventing mitochondrial Ca2+ overload.

## RESULTS

### TNF-mediated necrosis is dependent on cathepsin D, BID and BAX

Why might a necrosis pathway that could, in theory, be completed in a mitochondrion-intrinsic fashion (Baines, 2010; Bernardi, 2013; Halestrap, 2005; Nieminen, 2003; Zamzami et al., 2005) also require lysosomal components, i.e. lysosomal ceramide? A literature survey revealed a series of clues to this conundrum. Lyosomal ceramide is reported to mediate TNF-induced caspase-independent programmed cell death in a fibroblast cell line (Dumitru and Gulbins, 2006; Heinrich et al., 2004; Heinrich et al., 1999; Taniguchi et al., 2015; Thon et al., 2005). In vitro, ceramide produced by acid sphingomyelinase has been shown to activate lysosomal proteases, including cathepsins B and D (Heinrich et al., 2004; Heinrich et al., 1999; Taniguchi et al., 2015). In cell culture and in biochemical assays in vitro, cathepsins B and D have been shown to cleave the pro-apoptotic protein BID to its active form tBID (Appelqvist et al., 2012; Cirman et al., 2004; Droga-Mazovec et al., 2008; Heinrich et al., 2004). tBID activates BAX, a classical effector of apoptosis ((Karch et al., 2013; Karch et al., 2017; Westphal et al., 2011; Whelan et al., 2012), and the cancer chemotherapy drug gemcitabine was found to affect BAX-dependent apoptosis of a glioma cell line through a pathway that involves lysosomal accumulation of ceramide and cathepsin D activation (Dumitru and Gulbins, 2006; Heinrich et al., 2004; Heinrich et al., 1999; Taniguchi et al., 2015; Thon et al., 2005). More recently BAX is reported to work in conjunction with cyclophilin D to mediate mitochondrially-mediated necrosis (Karch et al., 2013; Karch et al., 2017; Westphal et al., 2011; Whelan et al., 2012). Collectively, these findings led us to consider the following sequence of events by which lysosomal components mediate necrosis: lysosomal ceramide activates cathepsin B and/or D which activate(s) BID and thereby BAX.

If this sequence is operant, then inhibiting cathepsin B and/or D, BID and BAX should inhibit TNF-mediated macrophage necrosis. We used bacterial cording as a read-out of macrophage necrosis; when macrophages undergo necrotic death, loss of granuloma cellularity renders bacteria extracellular (Figure 1B left panel); their exuberant growth in the permissive extracellular milieu is characterized by a readily-discernible serpentine cording morphology that can be quantified (Figure 1B right panel and Figure 1C) (Clay et al., 2008; Tobin et al., 2012). We inhibited cathepsins B and D, respectively, using the cysteine cathepsin inhibitor E64d and the aspartyl cathepsin inhibitor pepstatin A (Baldwin et al., 1993; Berg et al., 2016). Only pepstatin A inhibited TNF-mediated necrosis, pinpointing cathepsin D (Figure 1D and 1E). This was confirmed by genetic cathepsin D depletion using an antisense morpholino (Follo et al., 2011). Cathepsin D morphants were resistant to TNF necrosis (Figure 1F). Morpholino-mediated BID knockdown and pharmacological inhibition with BI-6C9, a small molecule tBID inhibitor (Becattini et al., 2004) also reduced necrosis (Figure 1G and 1H).

Next we asked if BAX was involved. The zebrafish BAX gene has undergone duplication so that there are two paralogs of human BAX, BAXA and BAXB, located adjacent to each other (Kratz et al., 2006). Sequence analysis revealed the highest values for identity, similarity and overall global/local score between human BAX transcript alpha and zebrafish BAXA, with conservation of important domains of the protein (Table S1 and Figure S1A). Although BAXB has been suggested to be the zebrafish functional equivalent of human BAK (Kratz et al., 2006), neither it nor BAXA had significant homology to either human BAK transcript (Table S1). We found that BAXA mutant zebrafish had substantial reduction in camptothecin-induced apoptosis, a BAX-dependent process in mammals (Albihn et al., 2007) (Figure S1B and C). BAXA was also required for TNF-mediated necrosis: BAXA morphants and mutants were resistant to it (Figure 1I and 1J). BAXB mutants had less reduction in camptothecin-induced apoptosis and were susceptible to TNF-mediated necrosis (Figure S1B and C and Figure 1K and 1L). Thus, in the zebrafish, BAXA has a greater role in apoptosis than BAXB and an exclusive role in TNF-mediated necrosis. In sum, our findings are consistent with lysosomal ceramide mediating necrosis through the activation of cathepsin D leading to BID and thereby BAXA (BAX) activation.

### BAX’s role in TNF-mediated necrosis is independent of oligomerization

BAX is known to participate in mitochondrially-mediated apoptosis through recruitment to and oligomerization on the mitochondrial outer membrane to form pores (Dewson and Kluck, 2009). More recently, BAX has been reported to also target the mitochondrial outer membrane to mediate necrosis in conjunction with cyclophilin D working in the inner mitochondrial membrane (Karch et al., 2015; Karch et al., 2013; Whelan et al., 2012). Different studies report this newly-identified role to be either dependent on or independent of its oligomerization function, which requires its BH3 domain (Karch et al., 2015; Karch et al., 2013) (Figure 2A). To test if BAX BH3 was required in TNF-mediated necrosis, we expressed BAX lacking the BH3 domain (ΔBH3-BAX) in BAX-deficient animals. We first confirmed that the BH3 domain was required for apoptosis as expected: while, as shown before (Kratz et al., 2006), expression of full length BAX in newly-fertilized BAX-deficient zebrafish eggs was rapidly lethal (due to exacerbated apoptosis), ΔBH3-BAX expression was not lethal (Figure S2A). BAX mutants expressing ΔBH3-BAX also remained resistant to camptothecin-mediated apoptosis (Figure S2C). In contrast, ΔBH3-BAX restored TNF-mediated necrosis, showing that BAX oligomerization and pore formation were not required (Figure 2B). Furthermore, BCB (BAX Channel Blocker), a small molecule inhibitor of BAX channel forming activity and cytochrome C release required for apoptosis (Cui et al., 2017), failed to inhibit macrophage necrosis, while inhibiting camptothecin-mediated apoptosis (Figure 2C and S2C). Together, these results showed that BAX functions in TNF-mediated necrosis independent of its oligomerization and pore forming activity.

**Figure 2:**
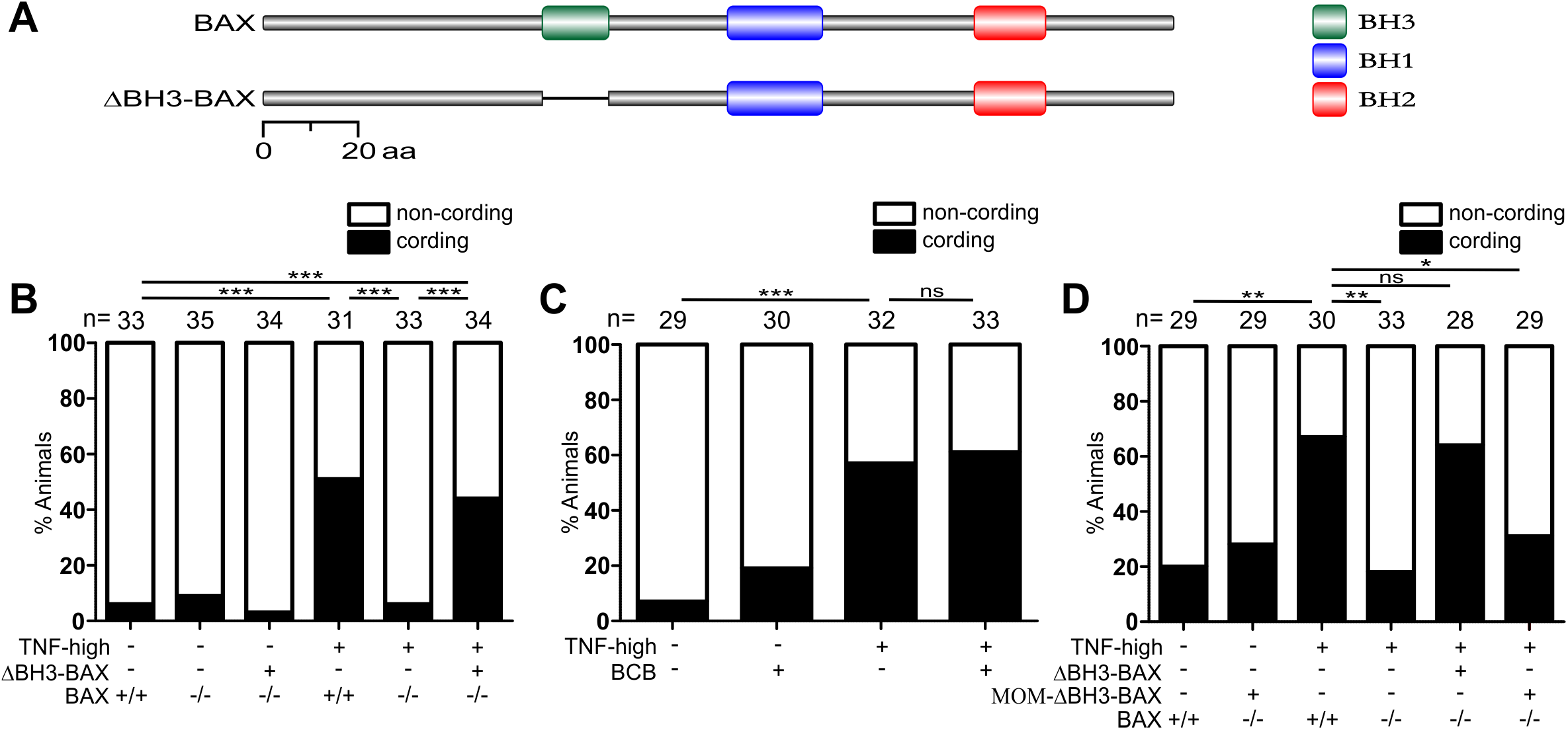
BAX mediates necrosis independent of both BH3-dependent oligomerization and its interaction with the mitochondrial outer membrane. (A) Schematic representation of BAX. Relevant BH domains of BAX are showed in colored boxes. (B) Percentage of animals with cording among WT and BAX mutants expressing ΔBH3-BAX injected with TNF or vehicle. ***p < 0.001 (Fisher’s exact test). Representative of 6 independent experiments. (C) Percentage of animals with cording among sibling larvae injected with TNF or vehicle in presence or absence of 5 μM BAX channel blocker (BCB). ***p < 0.001 (Fisher’s exact test). Representative of 3 independent experiments. (D) Percentage of animals with cording among WT and BAX mutant larvae expressing untagged or mitochondrial outer membrane-targeted ΔBH3-BAX injected with TNF or vehicle. *p < 0.05; **p < 0.01 (Fisher’s exact test). Representative of 2 independent experiments.

### BAX’s role in TNF-mediated necrosis does not require its localization to the outer mitochondrial membrane

Our findings so far were consistent with previous work using isolated mitochondria that indicated non-oligomerizing BAX interacts with the mitochondrion to mediate necrosis, working in conjunction with cyclophilin D (Karch et al., 2013; Whelan et al., 2012). If BAX was functioning thus in TNF-mediated necrosis, then expression of ΔBH3-BAX that is targeted to the mitochondrial outer membrane (Zhu et al., 1996) should be sufficient to restore macrophage necrosis in BAX-deficient animals. Strikingly, it did not restore necrosis (Figure 2D). We also confirmed that the mitochondrial tag did not disrupt BAX activity by showing that the apoptotic function of full length BAX (which is dependent on its proper mitochondrial insertion) bearing the mitochondrial tag was intact. Expression of this construct was similarly lethal to newly-fertilized BAX-deficient eggs as untagged BAX (Figure S2D). Together these results showed that BAX’s role in cyclophilin D-dependent necrosis is not through targeting the mitochondrial outer membrane.

### BAX mediates necrosis by promoting Ca2+ translocation from the ER to the mitochondrion

How then might BAX facilitate cyclophilin D-dependent necrosis? In addition to its mitochondrial interactions, BAX is reported to regulate calcium release from the ER, the major cellular Ca2+ repository (Kiviluoto et al., 2013; Nutt et al., 2002; Rong and Distelhorst, 2008; Rong et al., 2008; Rong et al., 2009; Vervliet et al., 2014; Vervliet et al., 2015; Vervliet et al., 2016). Because mitochondrial Ca2+ overload is a known trigger (in addition to oxidative stress) for the Cyclophilin D-dependent mPTP opening that causes necrosis (Clarke et al., 2002; Halestrap, 2005; Halestrap et al., 2004; Halestrap et al., 1998; Orrenius et al., 2003), we wondered if BAX worked in TNF-mediated necrosis by facilitating calcium flow from the ER to the mitochondrion to cause this overload. This model leads to five testable predictions 1) TNF-mediated necrosis should be associated with mitochondrial Ca2+ overload, 2) Inhibiting mitochondrial Ca2+ overload should rescue necrosis, 3) Mitochondrial Ca2+ overload should be BAX dependent as well as dependent on its upstream activators, ceramide, cathepsin D and BID, 4) BAX’s action in the ER should be sufficient to cause mitochondrial Ca2+ overload and necrosis and 5) Reducing ER Ca2+ stores should prevent necrosis.

To test the first prediction, that necrosis should be associated with mitochondrial Ca2+ overload, we used two approaches. First, we used mitochondrially-targeted GCaMP3 (a genetically-encoded Ca2+ indicator) to visualize mitochondrial Ca2+ (Esterberg et al., 2014). TNF increased the intensity of GCaMP3 fluorescence in macrophages of infected larvae within 90 minutes, indicating that it promoted mitochondrial Ca2+ uptake (Figure 3A). Consistent with only infected macrophages being susceptible to TNF-mediated necrosis, only infected macrophages in the TNF-high animals had increased mitochondrial Ca2+ (Figure 3B) (Roca and Ramakrishnan, 2013). In a complementary approach, we used Rhod-2, a fluorescent chemical probe for Ca2+ that accumulates in the mitochondrion (Smolina et al., 2014). Again infected, but not uninfected, macrophages became Rhod-2-positive within 120 minutes of TNF administration (Figure 3C and 3D). Moreover, mitochondrial Ca2+ overload by infected macrophages was rapidly followed by their necrosis; in contrast infected macrophages without Ca2+ overload did not die (Figure 3E). Thus, TNF caused mitochondrial Ca2+ overload selectively in infected macrophages and this mitochondrial Ca2+ overload preceded their necrosis.

**Figure 3.**
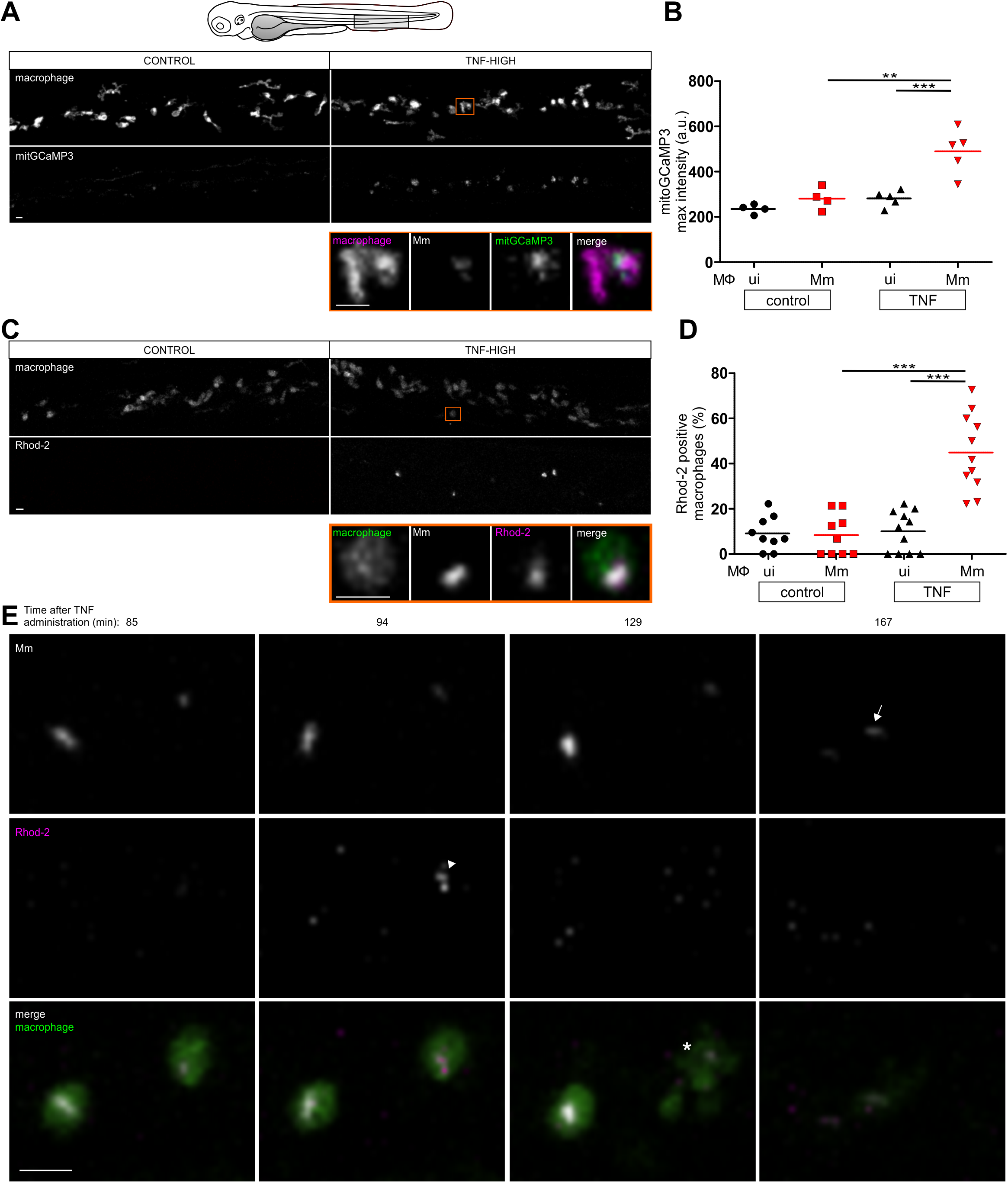
TNF promotes mitochondrial Ca2+ overload followed by macrophage necrosis. (A) Representative pseudo colored confocal images of 1 dpi *Tg(mfap4:mitGCaMP3)*;*Tg(mfap4:tdTomato-CAAX)* larvae 90 minutes after injection of TNF or vehicle corresponding to the shaded rectangle in the cartoon. Detail: macrophages in the orange rectangle. Scale bar 10 μm. (B) Quantitation of mitGCaMP3 fluorescence in individual macrophages from animals in (A). Each point represents the mean of mitGCaMP3 maximum intensity for uninfected or Mm-infected macrophages per fish. Black and red symbols represent uninfected and Mm-infected macrophages, respectively, in the same control or TNF-treated animal. ***p < 0.001 (one-way ANOVA with Tukey’s post-test). Representative of 2 independent experiments. (C) Representative pseudo colored confocal images of 1 dpi *Tg(mpeg1:YFP)*^*w200*^ larvae 2 hours after injection of TNF or vehicle in combination with Rhod-2 corresponding to similar area of the fish as in (A). Detail: macrophage in the orange rectangle. Scale bar 10 μm. (D) Percentage of uninfected or Mm-infected macrophages positive for Rhod-2 fluorescence (See Experimental Procedures) from fish in (D) over a course of 5 hours after TNF administration. Black and red symbols represent uninfected and Mm-infected macrophages, respectively, in the same control or TNF-treated animal. ***p < 0.001 (one-way ANOVA with Tukey’s post-test). Representative of 2 independent experiments. (E) Time lapse confocal images of two infected macrophages in a 1 dpi larva at the indicated time points after TNF administration. White arrowhead shows Rhod-2 positivity; asterisk shows macrophage death; white arrow shows extracellular bacteria. Scale bar 10 μm.

To test the second prediction, that inhibiting mitochondrial Ca2+ overload should rescue macrophage necrosis, we treated animals with Ru360, an inhibitor of the mitochondrial Ca2+ uniporter (MCU), which constitutes the major route of Ca2+ entry into the mitochondrial matrix (Zazueta et al., 1999). Ru360 prevented Ca2+ overload, and, importantly, also inhibited macrophage necrosis (Figure 4A and 4B). Thus, Ca2+ overload is a prerequisite for macrophage necrosis. To test the third prediction, that BAX and its upstream activators should be required for mitochondrial Ca2+ overload, we looked for TNF-induced mitochondrial Ca2+ overload in infected BAX deficient animals. It was absent (Figure 4C). Mitochondrial Ca2+ overload was also absent in ceramide-, cathepsin D- and BID-deficient animals (Figure 4D and 4E). These findings showed that BAX was required for mitochondrial Ca2+ overload. To test the fourth prediction that BAX causes mitochondrial Ca2+ overload solely by its action in the ER, we expressed ER-targeted ΔBH3-BAX in BAX-deficient animals (Zhu et al., 1996). ER-targeted ΔBH3-BAX restored TNF-mediated mitochondrial Ca2+ overload and macrophage necrosis in BAX-deficient animals, similar to untargeted ΔBH3-BAX (Figure 4F and 4G). To test the fifth and final prediction that reducing ER Ca2+ should prevent TNF-mediated necrosis, we used two approaches. We used the chemical thapsigargin to inhibit the activity of ER membrane-resident SERCA proteins that transfer Ca2+ from the cytosol into the ER lumen (Chemaly et al., 2018). Thapsigargin has been shown to deplete ER Ca2+ while increasing cytosolic Ca2+ (Inesi and Sagara, 1992; Iwasaki et al., 2015). Thapsigargin treatment of the animals reduced macrophage mitochondrial Ca2+ overload and reduced necrosis (Figure 4H and 4I). Second, we overexpressed the ER Ca2+ leak channels TMBIM 3 and 6 in the animals so as to reduce steady state ER Ca2+ concentrations (Lisak et al., 2015; Robinson et al., 2011; Xu et al., 2008). Overexpression of either TMBIM 3 or 6 inhibited both TNF-mediated mitochondrial Ca2+ overload and macrophage necrosis (Figure 4J and 4K). In sum, these findings revealed the ER, not the mitochondrion, to be the target of activated BAX in TNF-mediated necrosis of Mm-infected macrophages, and showed that BAX activation is required for Ca2+ translocation from the ER to the mitochondrion.

**Figure 4.**
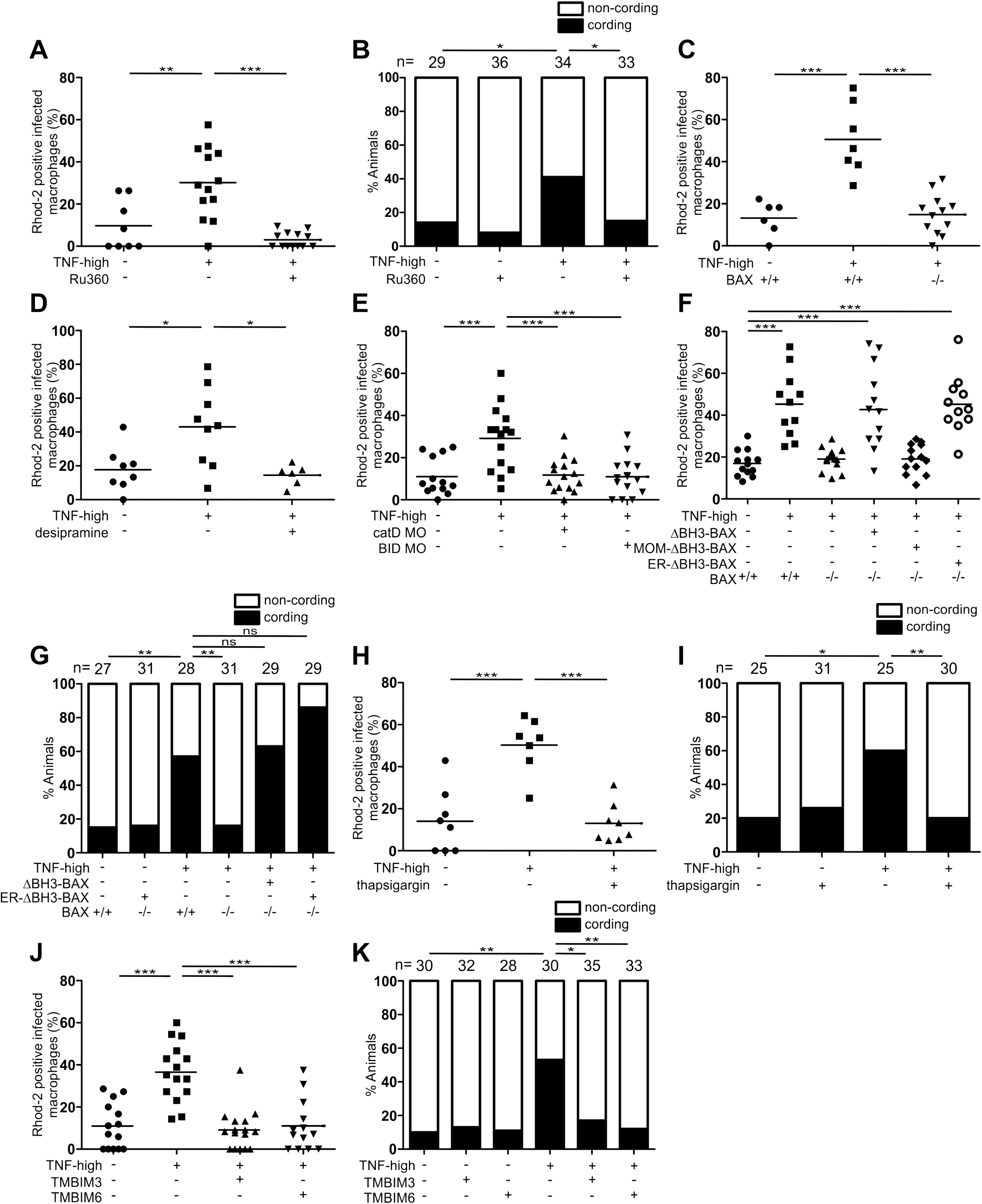
BAX promotes Ca2+ flow from the ER into the mitochondrion. (A) Percentage of Mm-infected macrophages which are positive for Rhod-2 fluorescence in 1dpi animals after TNF administration over a course of 5 hours in presence or absence of 2 μM Ru360. **p < 0.01; ***p < 0.001 (one-way ANOVA with Tukey’s post-test). Representative of 2 independent experiments. (B) Percentage of animals with cording among sibling larvae injected with TNF or vehicle in presence or absence of 2 μM Ru360. *p < 0.05 (Fisher’s exact test). Representative of 5 independent experiments. (C) Percentage of Mm-infected macrophages positive for Rhod-2 fluorescence in 1dpi animals among WT or BAX mutants after TNF administration over a course of 5 hours. ***p < 0.001 (one-way ANOVA with Tukey’s post-test). Representative of 2 independent experiments. (D) Percentage of Mm-infected macrophages positive for Rhod-2 fluorescence in 1dpi animals after TNF administration over a course of 5 hours in presence or absence of 10 μM desipramine. *p < 0.05 (one-way ANOVA with Tukey’s post-test). (E) Percentage of Mm-infected macrophages positive for Rhod-2 fluorescence in 1dpi WT and cathepsin D or BID morphants siblings after TNF administration over a course of 5 hours. ***p < 0.001 (one-way ANOVA with Tukey’s post-test). Representative of 2 independent experiments. (F) Percentage of Mm-infected macrophages positive for Rhod-2 fluorescence in 1dpi WT, BAX mutants and BAX mutant fish expressing untagged, MOM- or ER-targeted ΔBH3-BAX after TNF administration over a course of 5 hours. ***p < 0.001 (one-way ANOVA with Tukey’s post-test). Representative of 2 independent experiments. (G) Percentage of animals with cording among WT and BAX or BAX mutant larvae expressing untagged or ER-targeted ΔBH3-BAX injected with TNF or vehicle. **p < 0.01 (Fisher’s exact test). Representative of 2 independent experiments. (H) Percentage of Mm-infected macrophages which are positive for Rhod-2 fluorescence in 1dpi animals after TNF administration over a course of 5 hours in presence or absence of 1.5 μM thapsigargin. ***p < 0.001 (one-way ANOVA with Tukey’s post-test). (I) Percentage of animals with cording among sibling larvae injected with TNF or vehicle in presence or absence of 1.5 μM thapsigargin. *p < 0.05; **p < 0.01 (Fisher’s exact test). Representative of 3 independent experiments. (J) Percentage of Mm-infected macrophages positive for Rhod-2 fluorescence in 1dpi WT or TMBIM3- or TMBIM6-overexpressing fish after TNF administration over a course of 5 hours. ***p < 0.001 (one-way ANOVA with Tukey’s post-test). Representative of 2 independent experiments. (K) Percentage of animals with cording among WT or TMBIM3- or TMBIM6-overexpressing sibling larvae injected with TNF or vehicle. *p < 0.05; **p < 0.01 (Fisher’s exact test). Representative of 2 independent experiments.

### BAX mediates direct Ca2+ flow from the ER to the mitochondrion through Ryanodine receptors

In the context of apoptosis, BAX has been implicated in regulating ER Ca2+ levels (Bonneau et al., 2013; Scorrano et al., 2003; Vervliet et al., 2016). This action is indirect and is through sequestering BCL-2, an anti-apoptotic protein that can reduce ER Ca2+ through interactions with multiple Ca2+ channels (Bonneau et al., 2013; Scorrano et al., 2003; Vervliet et al., 2016). As BAX can only bind BCL-2 via the BH3 domain, our finding that ΔBH3-BAX could still mediate necrosis in BAX-deficient animals ruled out BCL-2 sequestration.

We searched for alternative mechanisms by which BAX acts at the ER to cause mitochondrial Ca2+ overload. Apoptosis in some contexts is associated with Ca2+ translocation from the mitochondrion to the ER through the IP3 receptor (IP3R) and IP3R-deficient T cells are resistant to it (Jayaraman and Marks, 1997; Vance, 2014). We wondered if BAX was promoting mitochondrial Ca2+ transit through the IP3R. To test this we took advantage of the finding that BCL2 binds and inhibits IP3R through its BH4 domain (Rong et al., 2009). Expression of the 28-amino acid BCL2 BH4 domain inhibited mitochondrial Ca2+ overload and necrosis, suggesting that IP3R could be involved (Figure 5A and 5B). To confirm this, we used xestospongin C, a highly specific IP3R inhibitor that has been shown to be active in the zebrafish to inhibit IP3R-mediated ER to mitochondrial Ca2+ flow (Esterberg et al., 2014; Gafni et al., 1997). Surprisingly, xestospongin C did not inhibit mitcohondrial Ca2+ overload and necrosis, suggesting that IP3R was not involved (Figure 5C and 5D). In reconciling the disparity between the xestospongin C and BCL2 BH4 effects, we realized that BCL2 BH4 also inhibits another group of specific ER Ca2+ channels, the Ryanodine receptors (RyR) (Vervliet et al., 2014; Vervliet et al., 2016). While well-known to be expressed in excitable cell types, two of the three RyR isoforms, RyR1 and RyR2 are also expressed in human monocytes/macrophages (there are no published data for RyR3) (Table S2). In the zebrafish too, RyR1 and RyR2 are highly expressed in monocytes, while RyR3 has a low level of expression (Table S2). To ask if RyR were involved in macrophage necrosis, we took advantage of the finding that only these receptors, and not IP3R, are inhibited by the BH4 domain of BCL-XL (Vervliet et al., 2015; Vervliet et al., 2016). BCL-XL BH4 inhibited mitochondrial Ca2+ overload and necrosis similar to BCL-2 BH4, implicating RyR (Figure 5A and 5E). We then tested ryanodine, a specific RyR inhibitor, and it too inhibited TNF-mediated mitochondrial Ca2+ overload and macrophage necrosis (Figure 5C and 5D). Another specific RyR inhibitor, dantrolene, that is an approved human drug also inhibited mitochondrial Ca2+ overload and necrosis (Table S2 and Figure 5F and 5G) (Zhao et al., 2001). Conversely, we asked if pharmacological activation of RyR with the small molecule 4-Chloro-*m*-cresol (4CmC) (Baur et al., 2000; Jacobson et al., 2006; Zorzato et al., 1993) restored TNF-mediated necrosis in BAX mutants. We first confirmed 4CmC activity in the zebrafish by showing that it increased fluorescence of the mitochondrial Ca2+ reporter GCaMP3 (Figure 5H). 4CmC did restore TNF-mediated necrosis in BAX mutants, and this was blocked by inhibiting mitochondrial Ca2+ uptake with Ru360 (Figure 5I). Thus, BAX acts at the ER through RyR to cause mitochondrial Ca2+ overload and necrosis.

**Figure 5.**
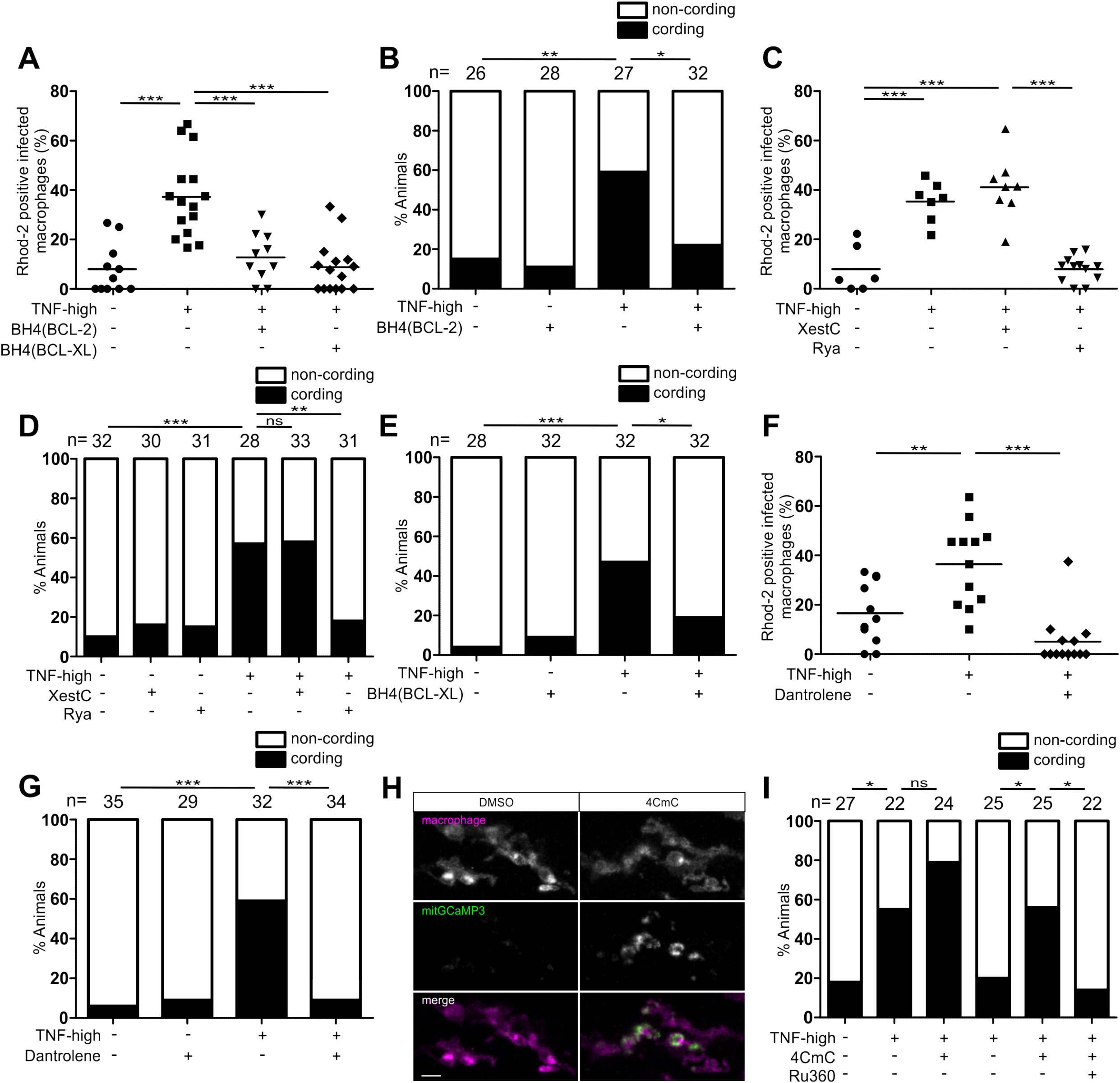
Ryanodine receptor mediates macrophage necrosis in excess TNF conditions. (A) Percentage of Mm-infected macrophages which are positive for Rhod-2 fluorescence in 1dpi WT or siblings expressing the BH4 domain of BCL-2 or BCL-XL after TNF administration over a course of 5 hours. ***p < 0.001 (one-way ANOVA with Tukey’s post-test). (B) Percentage of animals with cording among WT or siblings expressing the BH4 domain of BCL-2 injected with TNF or vehicle. *p < 0.05; **p < 0.01 (Fisher’s exact test). Representative of 3 independent experiments. (C) Percentage of Mm-infected macrophages which are positive for Rhod-2 fluorescence in 1dpi animals after TNF administration over a course of 5 hours in presence or absence of 5 μM Xestospongin C or 2 μM Ryanodine. ***p < 0.001 (one-way ANOVA with Tukey’s post-test). Representative of 2 independent experiments. (D) Percentage of animals with cording among sibling larvae injected with TNF or vehicle in presence or absence of 5 μM Xestospongin C or 2 μM Ryanodine. **p < 0.01; ***p < 0.001 (Fisher’s exact test). Representative of 2 independent experiments. (E) Percentage of animals with cording among WT or siblings expressing the BH4 domain of BCL-XL injected with TNF or vehicle. *p < 0.05; ***p < 0.001 (Fisher’s exact test). Representative of 2 independent experiments. (F) Percentage of Mm-infected macrophages which are positive for Rhod-2 fluorescence in 1dpi animals after TNF administration over a course of 5 hours in presence or absence of 5 μM dantrolene. **p < 0.01; ***p < 0.001 (one-way ANOVA with Tukey’s post-test). (G) Percentage of animals with cording among sibling larvae injected with TNF or vehicle in presence or absence of 5 μM dantrolene. ***p < 0.001 (Fisher’s exact test). Representative of 4 independent experiments. (H) Representative pseudo colored confocal images of 3 dpf *Tg(mfap4:mitGCaMP3)*;*Tg(mfap4:tdTomato-CAAX)* larvae 2 hours after administration of 10 μM 4CmC. Scale bar 10 μm. (I) Percentage of animals with cording among sibling larvae injected with TNF or vehicle in presence or absence of 10 μM 4CmC. *p < 0.05 (Fisher’s exact test). Representative of 2 independent experiments.

### TNF-induced mitochondrial ROS cause activation of lysosomal ASMase that ultimately results in mitochondrial Ca2+ overload

We now understood how the lysosomal pathway ultimately converged into the mitochondrion to cause cyclophilin D-dependent necrosis. One remaining question was how TNF triggers lysosomal ceramide production in infected macrophages. Because ASMase has been shown to be a redox sensitive enzyme, and mitochondrial ROS feature early in TNF-mediated necrosis (Roca and Ramakrishnan, 2013), we considered the possibility that mitochondrial ROS activate ASMase to produce ceramide. If so, then removing mitochondrial ROS should eliminate mitochondrial Ca2+ overload. We tested multiple ROS scavengers, including the mitochondrion-specific antioxidant MitoTEMPO, that were previously shown to rescue TNF-mediated necrosis (Roca and Ramakrishnan, 2013). All eliminated mitochondrial Ca2+ overload (Figure 6A). Thus, TNF-induced necrosis results from a single axis that initiates in the mitochondrion with ROS production, and traverses the lysosome, cytosol and ER to produce Ca2+ overload back in the mitochondrion to produce cyclophilin D-dependent necrosis. Consistent with this proposed order, ceramide, cathepsin D, BID and BAX, and RyR which we knew to be upstream of mitochondrial Ca2+ production (Figures 4C-4E and 5F) were found to be downstream of mitochondrial ROS: TNF-induced ROS were preserved in cathepsin D and BID morphants, BAX mutants, and in animals treated with the ASMase inhibitor desipramine and the RyR inhibitor dantrolene (Figure 6B-6D). As expected, mitochondrial ROS were not induced in animals treated with the RIPK1 inhibitor necrostatin-1 (Figure 6C), consistent with RIPK1 being upstream of ROS (Figure 1A)(Roca and Ramakrishnan, 2013). Finally, our model would predict that cyclophilin D mediates necrosis downstream of both ROS and Ca2+ production. To test this, we used cyclophilin D morphants and animals treated with the cyclophilin D inhibitor alisporivir. Cyclophilin D morphants and alisporivir-treated animals did not manifest TNF-induced macrophage necrosis, as predicted (Figure 6E)(Roca and Ramakrishnan, 2013). Yet mitochondrial ROS and Ca2+ were preserved in both groups of animals, confirming that cyclophilin D acts downstream of both signals (Figure 6F and 6G). Together these findings show that mitochondrial ROS initiate the circuit, and confirm the predicted order of the components.

**Figure 6.**
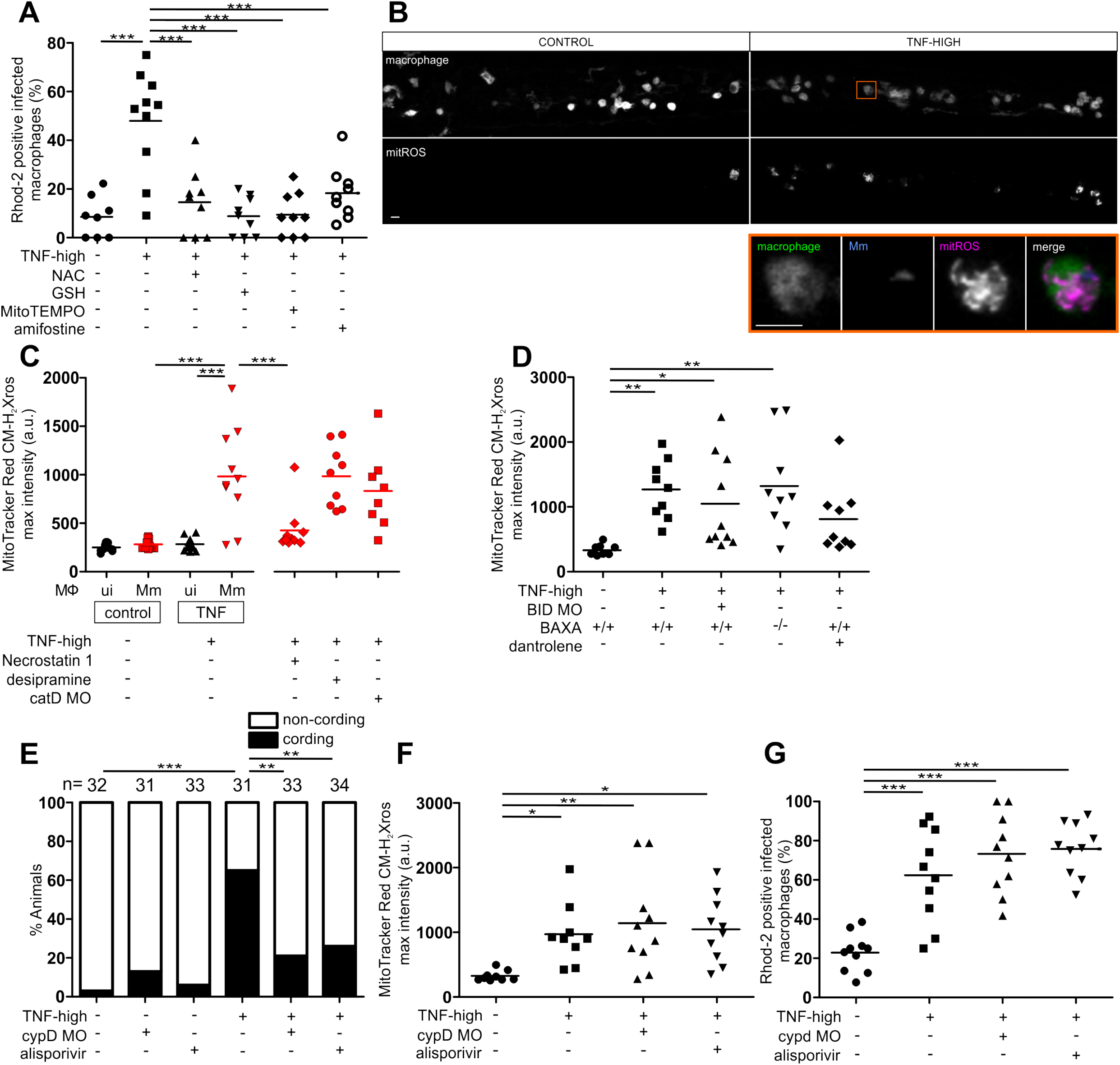
Mitochondrial ROS launch the pathogenic mitochondrial - lysosomal - ER circuit that leads to mitochondrial Ca2+ overload and macrophage necrosis. (A) Percentage of Mm-infected macrophages positive for Rhod-2 fluorescence in 1 dpi animals after TNF administration over a course of 5 hours in presence or absence of 40 μM NAC, 20 μM GSH, 20 μM mitotempo or 40 μM amifostine. ***p < 0.001 (one-way ANOVA with Tukey’s post-test). (B) Representative pseudo colored confocal images of 1 dpi *Tg(mpeg1:YFP)*^*w200*^ larvae 40 minutes after injection of TNF or vehicle in combination with MitoTracker Red CM-H_2_Xros corresponding to similar area of the fish as in Figure 3 (A). Detail: macrophage in the orange rectangle. Scale bar 10 μm. (C) Quantification of mitochondrial ROS production in 1 dpi control or cathepsin D morphant *Tg(mpeg1:YFP)*^*w200*^ larvae 40 minutes after injection of TNF or vehicle in combination with MitoTracker Red CM-H_2_Xros in presence or absence of 10 μM necrostatin-1 or 10 μM desipramine. Each point represents the mean of maximum intensity fluorescence of MitoTracker Red CM-H_2_Xros per fish from images in (B). Black and red symbols represent uninfected and Mm-infected macrophages, respectively, in the same control or TNF-administered animal. For necrostatin-1 and desipramine treated animals and for cathepsin D morphants, only Mm-infected macrophages were analyzed. ***p < 0.001 (one-way ANOVA with Tukey’s post-test). (D) Quantification of mitochondrial ROS production as in (C) in 1 dpi control or BID morphant or BAX mutant *Tg(mpeg1:YFP)*^*w200*^ larvae 40 minutes after injection of TNF or vehicle in combination with MitoTracker Red CM-H_2_Xros in presence or absence of 5 μM dantrolene. *p < 0.05; **p < 0.01 (one-way ANOVA with Tukey’s post-test). (E) Percentage of animals with cording among WT treated or not with 10 μM alisporivir and cyclophilin D morphant siblings injected with TNF or vehicle. **p < 0.01; ***p < 0.001 (Fisher’s exact test). (F) Quantification of mitochondrial ROS production in 1 dpi control or cyclophilin D morphant *Tg(mpeg1:YFP)*^*w200*^ larvae 40 minutes after injection of TNF or vehicle in combination with MitoTracker Red CM-H_2_Xros in presence or absence of 10 μM alisporivir. *p < 0.05; **p < 0.01 (one-way ANOVA with Tukey’s post-test). (G) Percentage of Mm-infected macrophages positive for Rhod-2 fluorescence in 1 dpi WT in presence or absence of 10 μM alisporivir or cyclophilin D morphant siblings after TNF administration over a course of 5 hours. ***p < 0.001 (Fisher’s exact test).

### Pharmacological inhibition of cellular Ca2+ uptake inhibits TNF-mediated mitochondrial Ca2+ overload and necrosis

Given the critical importance of mitochondrial Ca2+ overload for necrosis, we wondered if reduction of Ca2+ uptake into the macrophage would reduce steady state ER Ca2+ levels sufficiently so as to prevent mitochondrial Ca2+ overload. Nifedipine (dihydropyridine), diltiazem (benzothiazepine) and verapamil (phenylalkylamine) represent different classes of drugs that inhibit voltage gated L-type calcium channels (LTCCs) located in the plasma membrane (Ortner and Striessnig, 2016; Striessnig et al., 2015; Zamponi et al., 2015). While LTCCs have long been thought to be restricted to excitable cells (neurons, muscle cells and endocrine cells), recent work has found them to be expressed in a variety of cell types including immune cells (Davenport et al., 2015; Espinosa-Parrilla et al., 2015; Song et al., 2015; Zamponi et al., 2015). Importantly, LTCCs are present in in human, mouse and zebrafish myeloid cells; in humans these have been shown to include blood monocytes and alveolar macrophages where they have been shown to modulate immune and inflammatory functions such as macrophage activation and cytokine production (Antony et al., 2015; Athanasiadis et al., 2017; Ebina-Shibuya et al., 2017) (www.proteinatlas.org; www.immgen.org).

When we administered nifedipine, diltiazem and verapamil to TNF-high infected animals, mitochondrial Ca2+ overload and macrophage necrosis were inhibited (Figure 7A-7D). Thus, TNF-mediated necrosis could be inhibited by decreasing overall Ca2+ levels in macrophages through blocking LTCCs.

**Figure 7.**
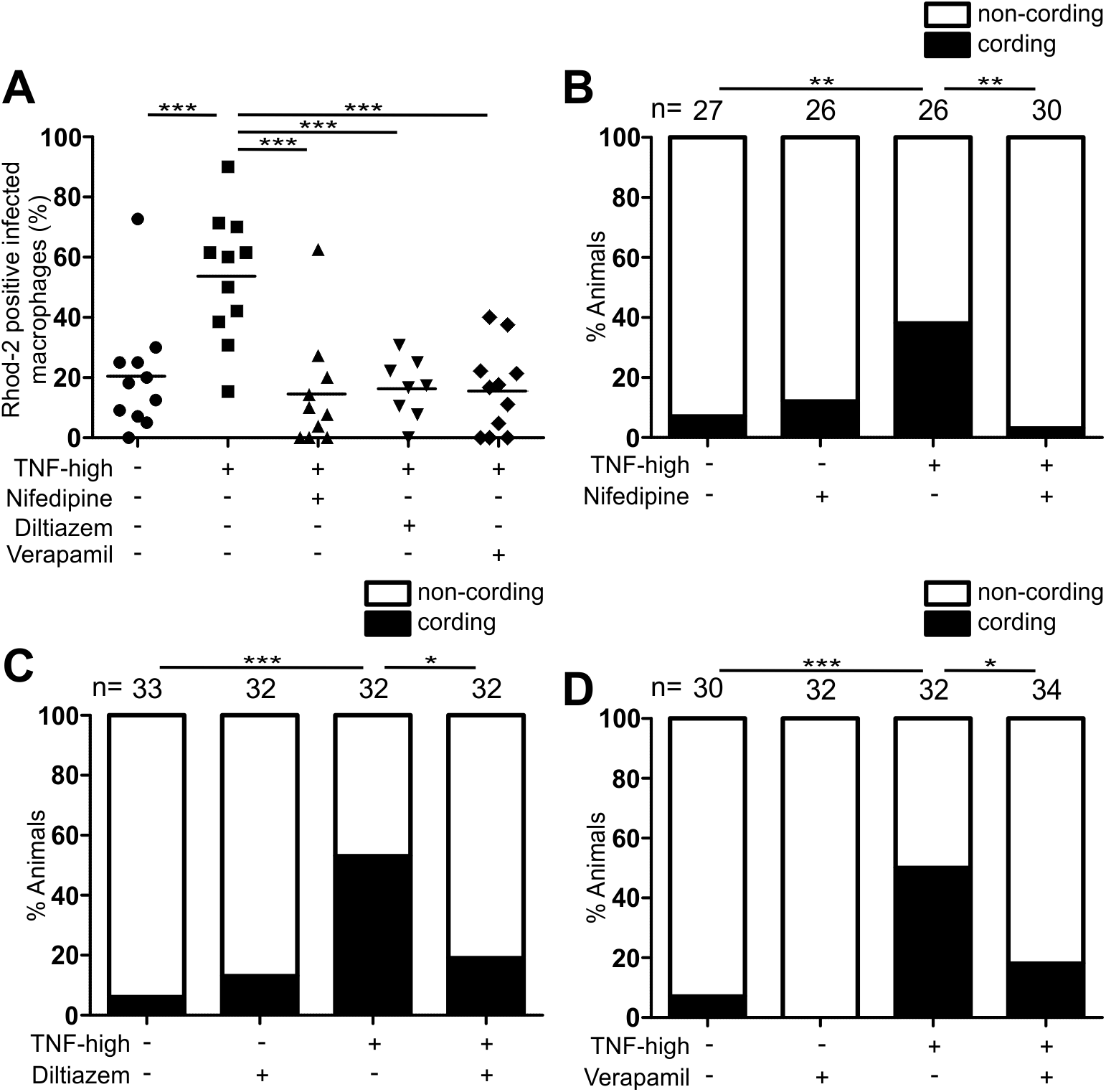
Pharmacological inhibition of voltage-gated L-Type Calcium channels inhibits necrosis. (A) Percentage of Mm-infected macrophages which are positive for Rhod-2 fluorescence in 1 dpi animals after TNF administration over a course of 5 hours in presence or absence of 3 μM nifedipine, 10 μM verapamil or 5 μM diltiazem. **p < 0.01; ***p < 0.001 (one-way ANOVA with Bonferroni’s post-test). Representative of 2 independent experiments. (B) Percentage of animals with cording among sibling larvae injected with TNF or vehicle in presence or absence of 3 μM nifedipine. **p < 0.01 (Fisher’s exact test). Representative of 4 independent experiments. (C) Percentage of animals with cording among larvae injected with TNF or vehicle in presence or absence of 5 μM diltiazem. *p < 0.05; ***p < 0.001 (Fisher’s exact test). Representative of 3 independent experiments. (D) Percentage of animals with cording 5dpi among larvae injected with TNF or vehicle in presence or absence of 10 μM verapamil. *p < 0.05; ***p < 0.001 (Fisher’s exact test). Representative of 2 independent experiments.

## DISCUSSION

Our dissection of the mechanism by which an excessive TNF response to mycobacterium infection causes host-detrimental necrosis of the infected macrophage has revealed an intricate interorganellar relay - a circuit that begins in the mitochondrion, transits the lysosome, cytosol and ER to complete in the mitochondrion (Figure 8). Our work assigns several pathway participants new roles, functions or links. We identify a new role for BAX as an activator of RyR. RyR are well-known Ca2+ translocators enabling the activity of excitable tissues like nerves and muscles. We find that these receptors are key mediators of pathology in mycobacterial infection through its activity in macrophages. Our dissection of the pathway reveals new interconnections between the participants that intriguingly span different organelles. The circuit begins and ends with the transit of two inorganic signals - ROS from mitochondrion to lysosome, and Ca2+ from ER to mitochondrion. It also requires cathepsin D translocation from lysosome to cytosol. The detailed understanding of the circuit suggests the potential of readily available, widely used drugs as host-targeting TB drugs.

**Figure 8.**
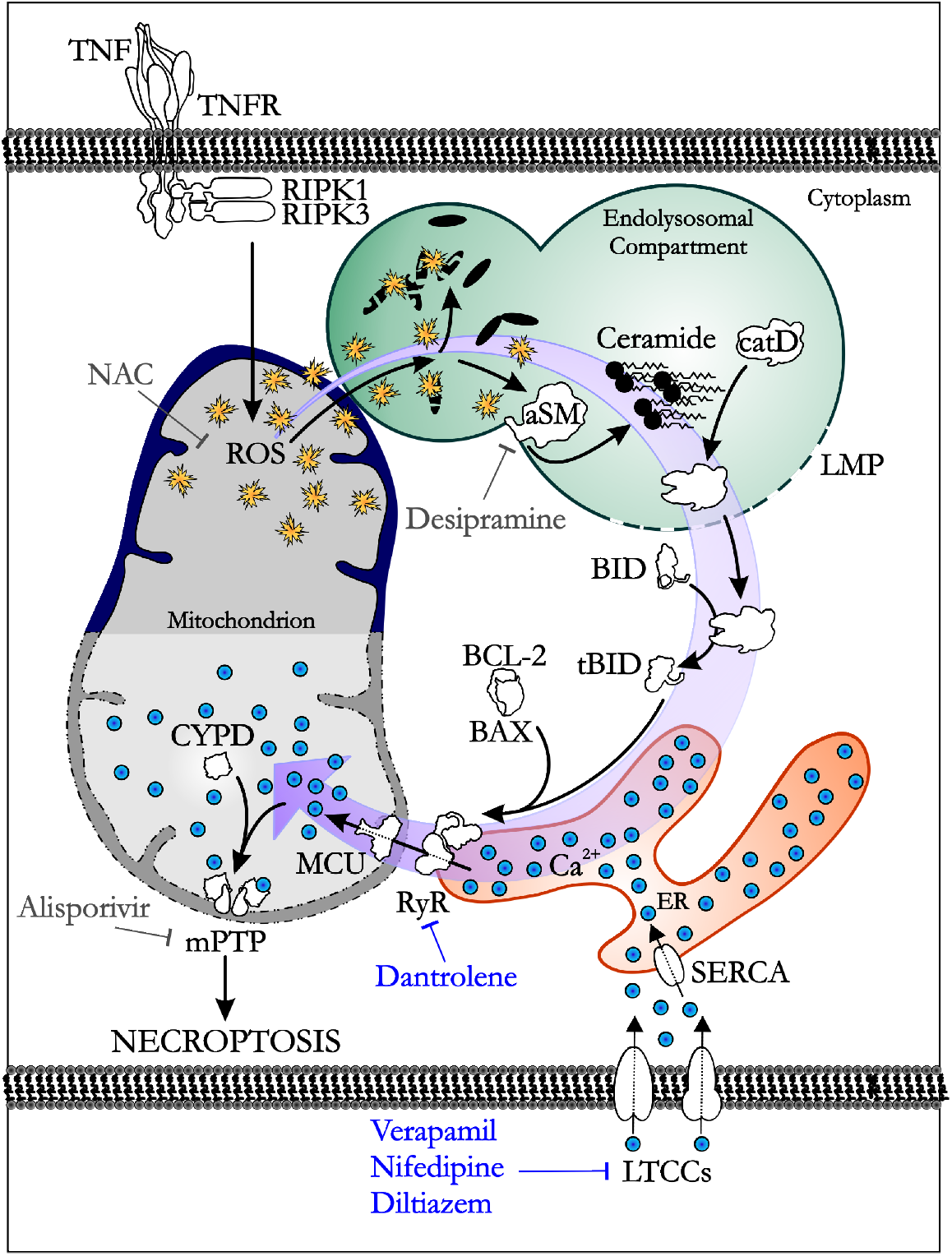
Model with circuit for TNF-mediated macrophage necrosis.

### Interorganellar contacts promote pathogenic macrophage necrosis

Ca2+ transfer between ER and mitochondrial channels is well documented and is facilitated by the formation of physical contacts between the two organelles - the so-called mitochondria-associated ER membranes (MAMs) (Naon and Scorrano, 2014; Thoudam et al., 2016; Vance, 2014). These tight connections allow for direct Ca2+ transit, creating a high Ca2+ concentration in the vicinity of the mitochondrial Ca2+ uniporter MCU located in the inner mitochondrial membrane, essential for Ca2+ transit through this low affinity receptor (Patron et al., 2013; Penna et al., 2018; Raffaello et al., 2012). Likewise, the RyR-MCU-mediated ER-mitochondrial Ca2+ transfer for TNF-mediated necrosis also appears to require direct contact; translocating Ca2+ from the ER to the cytosol by overexpressing ER Ca2+ leak channels removes mitochondrial Ca2+ overload and rescues necrosis. This is consistent with findings showing that RyR activity is coupled to VDAC, directly or indirectly interacting with it to promote Ca2+ translocation from the ER into mitochondrion (Fernandez-Sanz et al., 2014; Min et al., 2012).

In terms of the transit of ROS from mitochondrion to lysosome that initiates the necrosis pathway, recent studies find that mitochondrion - lysosome contacts also occur at appreciable frequency (Valm et al., 2017; Wong et al., 2018). This suggests a mechanism by which ROS, highly diffusible molecules, might be preferentially targeted to lysosomes. Importantly, these contacts form between healthy mitochondria and lysosomes, a finding germane to our pathway, where the transfer of mitochondrial ROS is an early event that must precede mitochondrial damage (Figure 8) (Wong et al., 2018). Furthermore, we have found that initially, prior to necrosis, these TNF-induced ROS are bactericidal to mycobacteria which reside in phagosomes and phagolysosomes, a function which may also be facilitated by targeted ROS delivery to these bacterium-containing compartments (Levitte et al., 2016; Roca and Ramakrishnan, 2013). Interorganellar contacts are thought to be part of cellular homeostasis and our findings suggest that they play critical roles in disease pathogenesis as well.

### Mitochondrial ROS translocate calcium into the mitochondrium through an elaborate interorganellar circuit

Mitochondrial oxidative stress is thought to trigger mitochondrial transition pore opening by oxidizing and regulating cyclophilin D activity (Baines et al., 2005; Du and Yan, 2010; Halestrap, 2005; Lopez-Erauskin et al., 2012; Martin, 2010). Therefore, prior to this study, it was reasonable to think that TNF-mediated mitochondrial ROS were mediating necrosis by directly activating cyclophilin D (Roca and Ramakrishnan, 2013). Instead, we now find that ROS activate lysosomal components that ultimately bring the Ca2+ into the mitochondrion that is essential for necrosis. Ca2+ is a well-known regulator of cyclophilin D activity and mitochondrial transition pore opening (Halestrap et al., 2004; Martin, 2010; Schinzel et al., 2005; Zoratti and Szabo, 1995). It is possible that mitochondrial ROS additionally work to directly regulate cyclophilin D in this pathway. However, this work shows that it is unlikely the predominant role as interventions that remove mitochondrial Ca2+ uptake while maintaining mitochondrial ROS production, inhibit necrosis almost completely. Thus, ROS, in the absence of Ca2+ are not sufficient to trigger cyclophilin D-dependent MPTP opening in this TNF-mediated necrosis. This work suggests these two inorganic signals co-operate to produce necrosis by each activating an enzyme - ROS the lysosomal enzyme ASMase, and Ca2+ the mitochondrial enzyme cyclophilin D.

### Extralysosomal cathepsin D function in TNF-mediated necrosis

The role assigned to cathepsin D in TNF-mediated necrosis - cleaving BID to its active state - raises the question of how a lysosomal protease might make its way to the cytosol? Lysosomal protease release from massive lysosomal disruption can cause necrosis (Turk et al., 2002). However, we can safely infer that cathepsin D works in TNF-mediated necrosis in the absence of massive lysosomal disruption - blocking the pathway immediately downstream of cathepsin D rescues necrosis. Thus, our findings are most consistent with a relatively specific translocation of cathepsin D. Both selective lysosomal membrane permeabilization and cytosolic translocation of cathepsin D have been observed in multiple studies (Ferri and Kroemer, 2001; Turk et al., 2002). Germane to our pathway, local increases in lysosomal ROS and sphingosine have been shown to produce lysoselective disruptions, and oxidative stress has been associated with cathepsin D cytosolic translocation (Ferri and Kroemer, 2001; Kagedal et al., 2001). We speculate that the mitochondrial ROS that reaches the lysosome may play the dual role of activating aSMase and permeabilizing the lysosomal membrane, that coordinate to promote cathepsin D activation and release into the cytosol. Furthermore, lysosomal ceramide is converted to sphingosine, so overproduction of ceramide would also increase lysosomal sphingosine, which could further promote lysosomal permeabilization.

In terms of cathepsin D’s cytosolic function has been documented in multiple contexts, and specific to ours, cathepsin D does cleave BID at neutral pH, which explains its retaining activity in the cytosol (Appelqvist et al., 2012). Finally, our data raise the question of why cathepsin D (and not cathepsin B) is specifically required in the pathway given that both cathepsins are activated by ceramide in vitro and both are translocated in response to oxidative stress to the cytosol where both can cleave BID (Appelqvist et al., 2012; Cirman et al., 2004; Droga-Mazovec et al., 2008; Heinrich et al., 2004; Heinrich et al., 1999; Kagedal et al., 2001; Taniguchi et al., 2015). One possible explanation is that “emergency” cytosolic protease inhibitors e.g., cystatins, can inhibit cysteine proteases like cathepsin B but no such inhibitors have yet been identified for the sole lysosomal aspartyl protease cathepsin D (Turk et al., 2002). Notably, earlier work on cathepsin D deficient mice already highlighted the importance of cathepsin D’s extralysosomal function: cathepsin D deficient mice were found to maintain normal bulk lysosomal proteolysis while undergoing widespread tissue destruction and early death (Saftig et al., 1995). Thus, the elaborate mechanisms that coalesce to enable cathepsin D cytosolic activity and translocation are rooted in the requirement for its homeostatic proteolytic function. Our work now highlights that these same mechanisms in overdrive can enable TB pathogenesis.

### BAX as an activator of RyR to cause Ca translocation in mycobacterium-infected macrophages

BAX, initially identified as a mediator of apoptosis, is more recently appreciated to play a role in other types of programmed cell death - necrosis and autophagy (Feldstein et al., 2006; Irrinki et al., 2011; Karch et al., 2015; Karch et al., 2013; Karch et al., 2017; Tischner et al., 2012; Westphal et al., 2011; Whelan et al., 2012; Zong et al., 2003). In most cases, its function is through disrupting organellar membranes - predominantly, the mitochondrion but also the ER and lysosome - generally through oligomerization on them to form pores or alternatively destabilizing membranes through other means (Feldstein et al., 2006; Irrinki et al., 2011; Karch et al., 2015; Karch et al., 2013; Karch et al., 2017; Tischner et al., 2012; Westphal et al., 2011; Whelan et al., 2012; Zong et al., 2003). Given how well studied BAX is, we were surprised to identify a completely new function for BAX where it activates RyR on the ER. Whether the action is direct or indirect remains to be determined but we know this newly identified moonlighting function occurs independently of its oligomerization and pore forming function.

The involvement of RyR rather than IP3R is yet another unexpected element. IP3R are ubiquitously present on all cells, trigger Ca2+ release in all cells; IP3R activity can promote apoptotic cell death by enabling mitochondrial Ca2+ overload through ER MAMs (Vervliet et al., 2016). We now find a similar role for RyR, which are mainly thought to translocate Ca2+ in the context of nerve and muscle excitation. RyR promote the rapid ER Ca2+ release that is required for efficient muscle contraction and nerve stimulation, suggesting their highly specialized function (Lanner et al., 2010). Yet their expression in macrophages across species suggests that they must mediate Ca2+ transfer in nonexcitable cells as well, perhaps with different kinetics. Our findings not only highlight this function, albeit in a pathological context, but use its involvement to curb the process through the use of the well known RyR inhibiting drug, dantrolene.

### Dysregulated TNF as a general cause of tuberculous granuloma necrosis

Our quest to understand the detailed mechanism of TNF-mediated macrophage necrosis in the zebrafish was instigated by its potential relevance for TB severity and treatment in humans with a common genetic LTA4H variant that produces a hyperinflammatory state (Thuong et al., 2017; Tobin et al., 2012). However, the TNF-initiated circuit we have identified may well play a wider role in tuberculous granuloma necrosis. Macrophages produce TNF in response to virulent mycobacteria even in the absence of high LTA4H. This TNF is a critical protective determinant, and only its excess launches the circuit described here (Clay et al., 2008; Roca and Ramakrishnan, 2013; Tobin et al., 2012). While excess LTA4H activity is one cause of excess TNF production, dysregulated TNF is probably a more general feature in TB, driven by multiple mechanisms. Human macrophages can undergo necrosis when infected with virulent Mtb, and a recent study finds that necrosis of Mtb-infected human macrophages is dependent on RIPK3, mitochondrial ROS and cyclophilin D, suggesting that the same circuit we describe is in play (Zhao et al., 2017). We wonder if TNF is the driver of necrosis in those studies too, with relatively high macrophage bacterial burdens tipping TNF production over the threshold from beneficial to excessive.

Likewise, the necrosis seen in nearly all advanced human TB granulomas may result from local TNF excess. A minority of macrophages may undergo necrosis for a variety of reasons including bacterial overload due to reduced microbicidal capacity (Ramakrishnan, 2012). Macrophages undergoing necrosis release the danger signal HMGB1, a potent TNF inducer that would stimulate TNF production in neighboring live macrophages, thus extending necrosis through the TNF pathway (Kaczmarek et al., 2013). Therefore the TNF-mediated necrosis pathway could be operant in the general context of TB granulomas, even independent of LTA4H genotype. In support of this idea, lactosylceramide, a downstream product of ceramide that is increased in cells by pro-inflammatory cytokines, has also been found to be increased in human lung necrotic granulomas (Chatterjee and Pandey, 2008; Kim et al., 2010). This lactosylceramide may, in fact, be the result of further metabolism of acid sphingomyelinase-derived ceramide in granuloma macrophages undergoing TNF-mediated necrosis. Thus, TNF mediated necrosis may be a more general feature of TB, and the high LTA4H genotype or other genetically mediated dysregulated TNF responses may only serve to increased susceptibility by jumpstarting necrosis early in infection when macrophage bacterial burdens are still low.

Our attempt to dissect TB pathogenesis has revealed new cell biology or at least new signaling circuitry. Many inflammatory conditions feature programmed necrosis that is increasingly appreciated to be detrimental in a wide range of noninfectious inflammatory pathologies (Zhou and Yuan, 2014) It is conceivable that some of these are the result of the pathway we have identified, initiated by dysregulated TNF coupled to a second nonbacterial trigger to induce mitochondrial ROS. Finally, the identification of mitochondrial calcium overload as a requisite for necrosis led us to ask if drugs block LTCCs would reduce calcium entry into the macrophage itself, and thereby prevent the pathological mitochondrial overload required for necrosis. Our finding that a panel of widely-used calcium channel blocking drugs prevent both macrophage mitochondrial Ca2+ overload and necrosis in TNF-high zebrafish suggest a potential for these drugs for TB sufferers with LTA4H-high genotype, and possibly more generally for TB as well as other inflammatory necroses where this pathway might be operant.

## AUTHOR CONTRIBUTIONS

All authors conceived and designed experiments, FR and SR performed experiments, FR prepared figures, FR and LR wrote the paper.

## ACKNOWLEDGMENTS

We thank D. Green, J. Prudent, A. Whitworth for advice and discussion, D. Raible for providing plasmids containing GCaMP3, Novartis for providing alisporivir, N. Goodwin, R. Keeble, and J. Cameron for zebrafish husbandry, A. Almeida, N. Goodwin, and R. Keeble for genotyping mutants, A. Almeida for technical assistance in creating the mfap4:GCaMP3 line, and L. Whitworth for manuscript review. This work was supported by an NIH R01 grant and a Wellcome Trust Principal Research Fellowship to LR.

---

**Supplemental Figure 1 (related to Figure 1).**
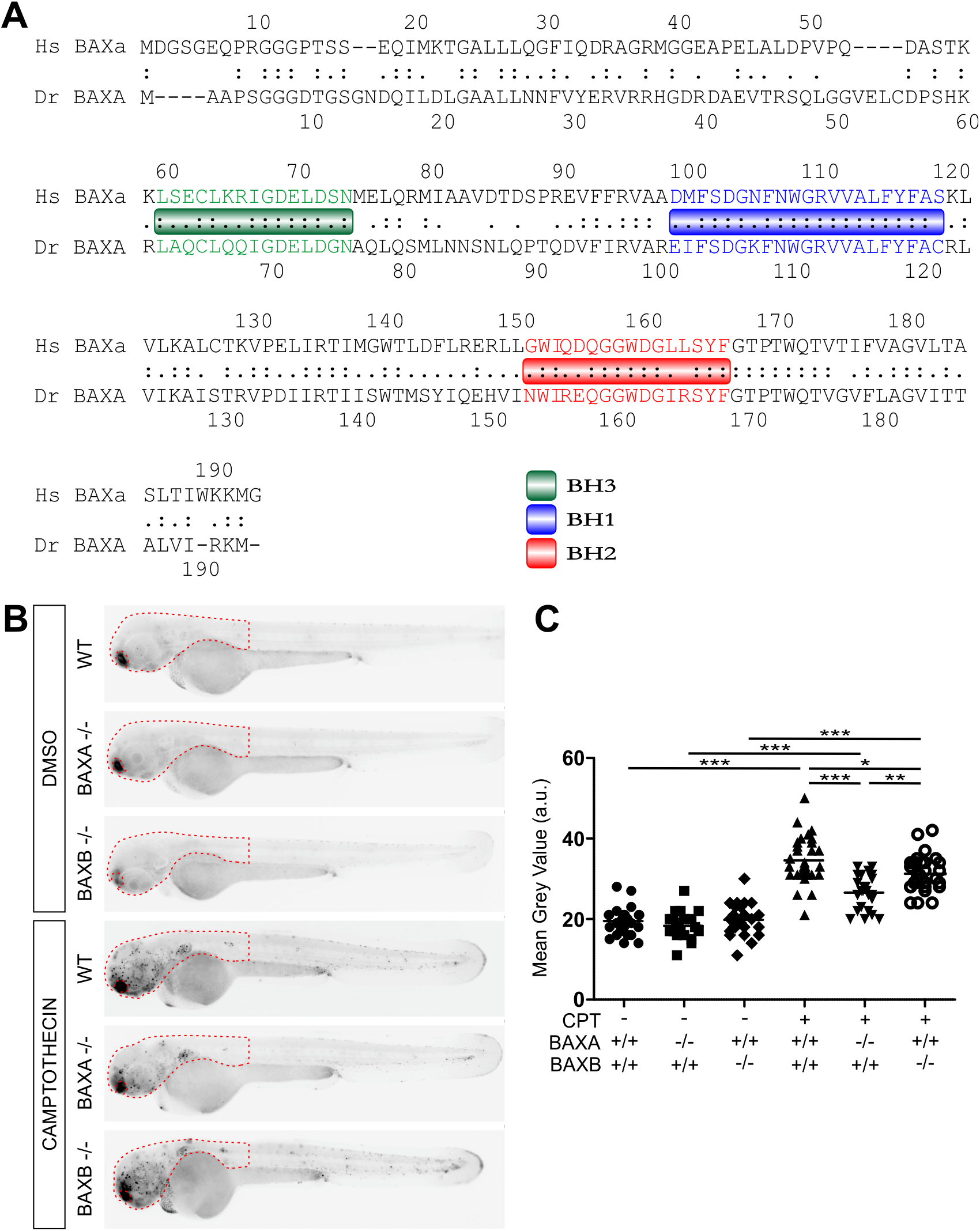
Camptothecin-induced apoptosis is decreased in BAXA and BAXB mutant zebrafish. (A) Comparison of protein sequence homology between human BAX alpha and zebrafish BAXA. Relevant BH domains of BAX are showed in colored boxes. (B) Representative inverted fluorescence images 2 dpf WT, BAXA mutant or BAXB mutant larvae in presence or absence of 500 nM camptothecin and incubated with acridine orange to detect apoptotic cells (See Experimental Procedures). (C) Quantification of camptothecin-induced apoptosis (See Experimental Procedures) in fish from (A) in the area delimited with red dashed line as showed in (A) in WT, BAXA mutant or BAXB mutant larvae. *p < 0.05; **p < 0.01; ***p < 0.001 (one-way ANOVA with Tukey’s post-test). Representative of 3 independent experiments.

**Supplemental Figure 2 (related to Figure 2).**
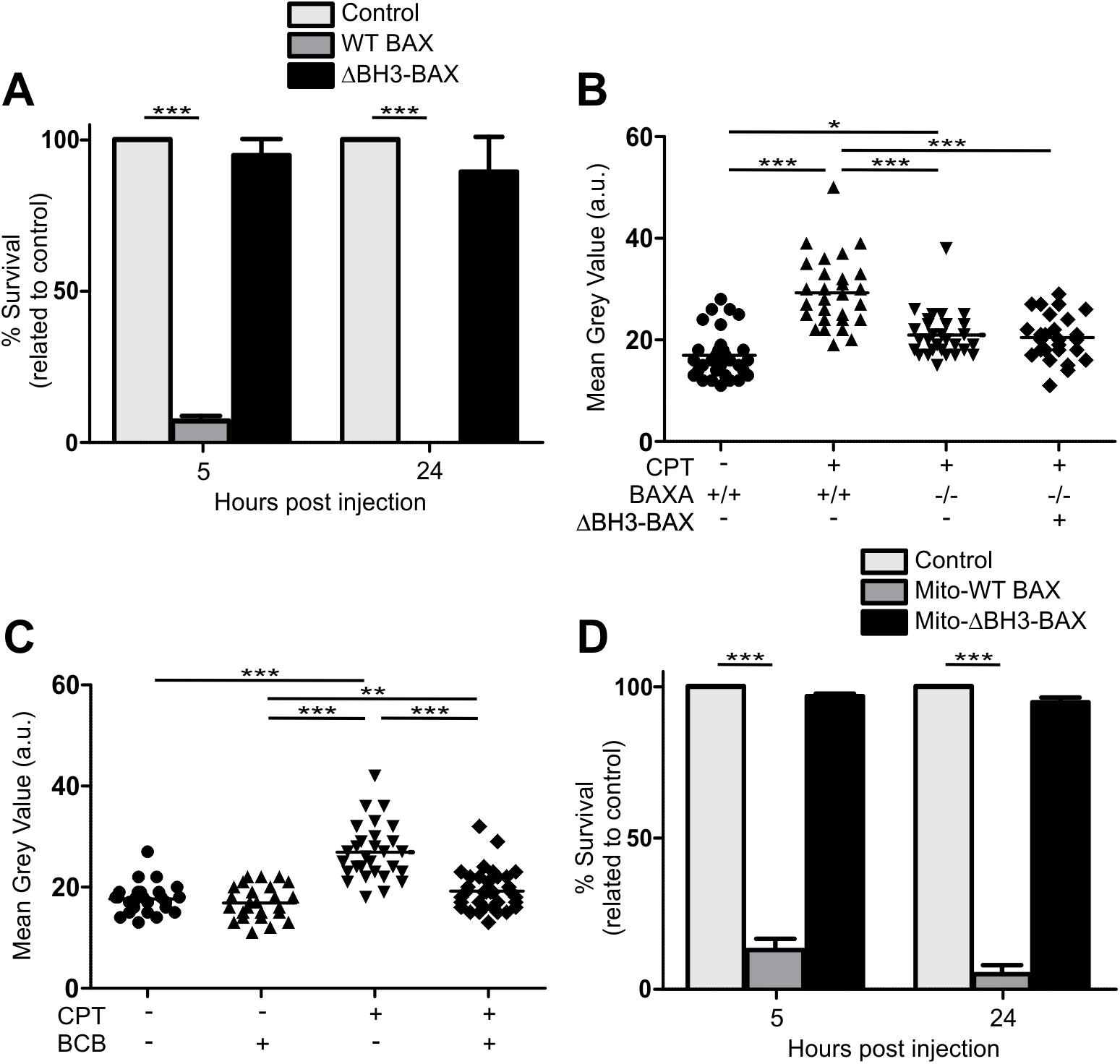
BH3 domain of BAX is required for apoptosis. (A) Percentage survival of BAX mutant embryos expressing WT or ΔBH3-BAX 5 and 24 hpf. ***p < 0.001 (one-way ANOVA with Tukey’s post-test). Mean of 3 experiments (±SEM) is represented. (B) Quantification of camptothecin-induced apoptosis in 2 dpf larvae among WT and BAXA mutants expressing or not ΔBH3-BAX. *p < 0.05; ***p < 0.001 (one-way ANOVA with Tukey’s post-test). Representative of 2 independent experiments. (C) Quantification of camptothecin-induced apoptosis in 2 dpf animals in presence or absence of 5 uM BCB. **p < 0.01; ***p < 0.001 (one-way ANOVA with Tukey’s post-test). Representative of 2 independent experiments. (E) Percentage survival of 5 and 24 hpf BAX mutant embryos expressing MOM-WT or MOM-ΔBH3-BAX. ***p < 0.001 (one-way ANOVA with Tukey’s post-test). Mean of 3 experiments (±SEM) is represented.

**Table S1.** Global analysis without end-gap penalty of protein sequence homology between human BAK1 (transcripts 202 and 203), BAX (only major transcripts, alpha and beta) and zebrafish BAXA and BAXB. Identity (%), similarity (%), (global/local score). Prefix h indicates human; prefix zf indicates zebrafish.

|  | **hBAXb** | **hBAK1-202** | **hBAK1-203** | **zfBAXA** | **zfBAXB** |
| --- | --- | --- | --- | --- | --- |
| **hBAXa** | **85.4**<br>88.5<br>(1056) | **22.3**<br>58.8<br>(183) | **20.8**<br>49.4<br>(36) | **49.6**<br>77.3<br>(631) | **23.1**<br>55.7<br>(212) |
| **hBAXb** |  | **21.9**<br>50.2<br>(126) | **20.8**<br>49.4<br>(36) | **36.6**<br>63.4<br>(456) | **22.9**<br>52<br>(193) |
| **hBAK1-202** |  |  | **78.4**<br>86.3<br>(754) | **22.4**<br>57.9<br>(154) | **24.8**<br>53<br>(126) |
| **hBAK1-203** |  |  |  | **19.9**<br>49.4<br>(13) | **21.9**<br>45.2<br>(4) |
| **zfBAXA** |  |  |  |  | **22.3**<br>59.1<br>(203) |

**Table S2.**
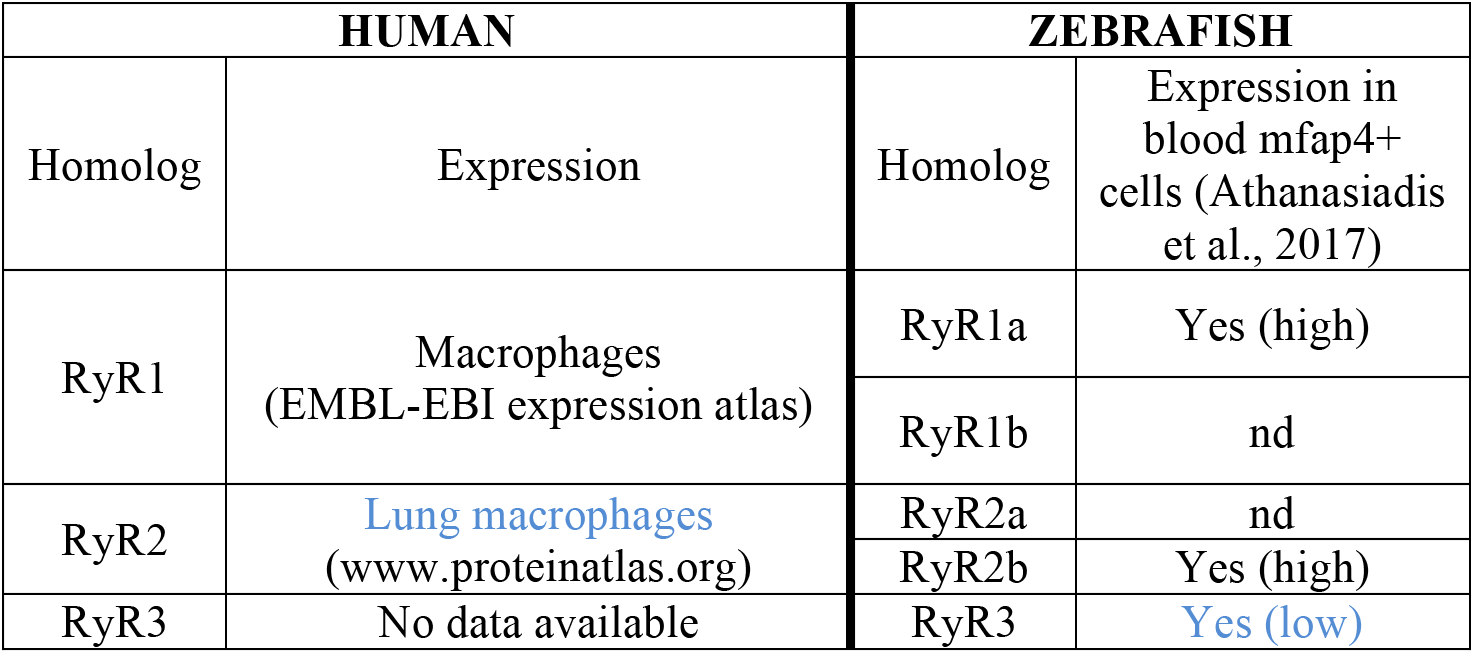
Presence of ryanodine receptor isoforms in human macrophages and zebrafish blood monocytes.

| HUMAN |  | ZEBRAFISH |  |
| --- | --- | --- | --- |
| Homolog | Expression | Homolog | Expression in blood mfap4+ cells (Athanasiadis et al., 2017) |
| RyR1 | Macrophages (EMBL-EBI expression atlas) | RyR1a | Yes (high) |
|  |  | RyR1b | nd |
| RyR2 | [Lung macrophages](http://www.proteinatlas.org) (www.proteinatlas.org) | RyR2a | nd |
|  |  | RyR2b | Yes (high) |
| RyR3 | No data available | RyR3 | Yes (low) |

## STAR METHODS

### CONTACT FOR REAGENT AND RESOURCE SHARING

Further information and requests for resources and reagents should be directed to and will be fulfilled by the Lead Contact, Lalita Ramakrishnan.

### EXPERIMENTAL MODEL AND SUBJECT DETAILS

#### Zebrafish husbandry and infections

Zebrafish husbandry and experiments were conducted in compliance with guidelines from the UK Home Office (experiments conducted in the University of Cambridge) and with the U.S. Public Health Service Policy on Humane Care and Use of Laboratory Animals using protocols approved by the Institutional Animal Care and Use Committee of the University of Washington (generation of Baxa^rr1^ and Baxb^rr10^ mutant lines in the University of Washington). Zebrafish AB wild-type strain (Zebrafish International Resource Center) and transgenic or mutant lines in the AB background were used, including Tg(mpeg1:YFP)^w200^ (Roca and Ramakrishnan, 2013), Tg(mpeg1:Brainbow)^w201^ (expressing tdTomato) (Pagan et al., 2015), Tg(BH:GFP-mfap4:Mito-GCaMP3) (this work), baxa^rr1^ (this work) and baxb^rr10^ (this work). All zebrafish lines were maintained in buffered reverse osmotic water systems and were exposed to a 14 hr light - 10 hr dark cycle to maintain proper circadian conditions. Fish were fed twice daily a combination of dry food and brine shrimp. Zebrafish embryos were housed in fish water (reverse osmosis water containing 0.18 g/l Instant Ocean) at 28.5°C. Embryos were maintained in 0.25 μg/ml methylene blue from collection to 1 dpf. 0.003% PTU (1-phenyl-2-thiourea, Sigma) was added from 24 hours post-fertilization (hpf) on to prevent pigmentation. Larvae (of undetermined sex given the early developmental stages used) were anesthetized in fish water containing 0.025% tricaine (Sigma) and infected at 48 hpf via caudal vein (CV) injection using single-cell suspensions of Mm of known titer (Takaki et al., 2012; Takaki et al., 2013). Larvae were randomly allotted to the different experimental conditions. Number of animals to be used for each experiment was guided by pilot experiments to reach statistical significance and each experiment was repeated two or more times.

Generation of the transgenic zebrafish line Tg(BH:GFP-mfap4:Mito-GCaMP3): the plasmid pTol2-PhiC31LS-BH:GFP-mfap4:Mito-GCaMP3 was generated by PCR amplifying the mito-GCaMP3 cassette from the plasmid pME mitoGCaMP3 (Esterberg et al., 2014) and cloning into a Tol2 plasmid with a green bleeding heart cassette (cmlc2:eGFP) for screening [pTol2 PhiC31LS BH NewMCS (cmlc2:eGFP)] followed by cloning of the mfap4 promoter (Walton et al., 2015). The plasmid pTol2-PhiC31LS-BH NewMCS (cmlc2:RFP) was generated by NEBuilder HiFi DNA Assembly following the manufacturer protocol by combining the backbone of the Tol2 plasmid pTol2 mfap4:TdTomato-CAAX (Walton et al., 2015) with sequence containing the bleeding heart and Phi31 Landing cassettes from the plasmid pSB_PhiC31LandingSite (Kirchmaier et al., 2013) (Addgene plasmid 48875) where HA tag was previously removed and a new multi-cloning site inserted. A version of pTol2 PhiC31LS BH NewMCS with eGFP expression in the myocardium [pTol2 PhiC31LS BH NewMCS (cmlc2:eGFP)] was generated by replacing RFP with eGFP by NEBuilder HiFi DNA Assembly. The pTol2-PhiC31LS-BH:GFP-mfap4:Mito-GCaMP3 plasmid was injected along with transposase mRNA into one- to two-cell-stage embryos of the wild-type AB strain as previously described (Suster et al., 2011) using injection mix (1× Tango Buffer (Thermo Scientific) containing 2% phenol red sodium salt solution (Sigma)). Putative founders were identified by GFP expression in the heart and crossed to wild-type AB zebrafish. Transgenic lines were identified in the next generation and kept on the AB strain.

BAXA- and BAXB-deficient zebrafish lines were generated by using the CRISPR/Cas9 system (Irion et al., 2014) in the University of Washington. The mutation consists in a single nucleotide (G) deletion in exon 2 of baxa and a deletion of three nucleotides (TGG) and insertion of thirteen (AATAAAGAGGTGA) in exon 3 of baxb. Both Baxa^rr1^ and Baxb^rr10^ lines were genotyped by high-resolution melt analysis (HRM) (Garritano et al., 2009) of PCR products (see Key Resources Table for sequences) on a CFX Connect thermocycler (BioRad).

### METHOD DETAILS

#### Bacterial strains

*M. marinum* M strain (ATCC #BAA-535) expressing tdTomato or EBFP2 under control of the msp12 promoter (Takaki et al., 2013), were grown under hygromycin B (Formedium) selection in 7H9 Middlebrook medium (Difco) supplemented with oleic acid, albumin, dextrose, and Tween-80 (Sigma) (Takaki et al., 2013).

#### TNF and drug administration

0.5 ng of recombinant zebrafish soluble TNF (Roca et al., 2008) or vehicle were microinjected into the caudal vein of each animal 1 day post infection (Tobin et al., 2012). To assess drug treatment in infected fish, equivalently-infected sibling larvae were mixed in a petri dish and held at 28.5°C until they were then randomly allocated to the drug-treated or control. DMSO (Sigma) was kept at 0.5% in all conditions when drugs were being used. All drugs were dissolved in DMSO (Sigma) or water and kept in small aliquots at −20°C. The rational for the doses used in this work was based in previous studies or in pilot experiments, using the minimum effective concentration. None of these concentrations showed toxic effects in the animals at the end of the experiment. All drugs were administered by adding them to the fish water. BI-6C9 (5 μM) (Sigma), BCB (Bax Channel Blocker) (10 μM) (Alfa Aesar) and Alisporivir (10 μM) (Novartis) were administered 4 hours after infection and removed 24 hours post TNF administration. Pepstatin A (7.5 μM) (Sigma), E64d (1 μg/ml) (Sigma), Ru360 (2 μM) (VWR International), thapsigargin (1.5 μM) (Sigma), Amifostine (40 μM) (Cambridge Bioscience), NAC (30 μM) (Cambridge Bioscience), Mitotempo (20 μM) (Cambridge Bioscience), GSH (20 μM) (Cambridge Bioscience), Xestospongin C (5 μM) (Cambridge Bioscience), Ryanodine (2 μM) (Generon), Dantrolene (5 μM) (Sigma), 4-Chloro-*m*-cresol (4CmC) (10 μM) (Sigma), Nifedipine (3 μM) (Cambridge Bioscience), Diltiazem (5 μM) (Cambridge Bioscience) and Verapamil (10 μM) (Fischer Scientific) were administered 5 hours prior and removed 24 hours post TNF administration. For experiments quantifying mitochondrial Ca2+ overload, drugs were administered 5 hours prior to TNF administration and maintained during imaging.

#### Morpholino knockdown

Cathepsin D-splice-blocking and BID-, BAXA-, and cyclophilin D-translation-blocking morpholinos (see Key Resources Table for sequences) (Gene Tools) were diluted in injection solution and approximately 2-4 nl were injected into the yolk of one- to two-cell-stage embryos (Tobin et al., 2012). The concentration in the injection solution for each morpholino was 0.2 mM for Cathepsin D and 0.15 mM for BID, BAXA and cyclophilin D.

#### Synthetic mRNA synthesis and microinjection

The sequences for the ORFs for acid ceramidase, BAXA, BC-L2 and TMBIM3 and 6 were obtained by PCR from zebrafish cDNA. The ORF for acid ceramidase was then cloned into the plasmid pCS2P+ and mRNA was generated with T7 (Roca and Ramakrishnan, 2013). The T7 promoter followed by the Kozac sequence 5’-GCCGCCACC-3’ were inserted before the start codon by PCR for all versions of BAXA, Venus-2A-BH4 (from BC-L2) and TMBIM3 and 6. ΔBH3-BAX was generated by PCR by removing the BH3 domain. MOM-BAX and MOM-ΔBH3-BAX were generated by PCR by removing the Stop codon of BAX and fusing a linker sequence G_4_SG_4_SG_4_ followed by the MOM-targeting sequence from the protein ActA from Listeria monocytogenes (Zhu et al., 1996) and a Stop codon using as a template full length or ΔBH3-BAX, respectively. ER-ΔBH3-BAX was generated by PCR by removing the Stop codon of BAX and fusing the linker sequence G_4_SG_4_SG_4_ followed by the ER-targeting sequence from the rat protein cytochrome b5 (Zhu et al., 1996) and a Stop codon using as a template ΔBH3-BAX. Venus-2A-BH4 (from BC-L2) was generated by PCR. For larval survival experiments, the sequences for BAX and ΔBH3-BAX were cloned in the plasmid pCS2P+ downstream and in frame with the sequence for venus-V2A to ensure expression of the constructs. RNA was synthesized using the mMessage mMachine kit (Ambion) and the polyA Tailing kit (Ambion). 2-4 nl were injected into the yolk of one- to two-cell-stage embryos at different concentrations in injection solution.

#### Microscopy

Fluorescence microscopy was performed as described (Takaki et al., 2013). Quantification of camptothecin-induced apoptosis by acridine orange staining and assessments of mycobacterial cording were performed with a Nikon Eclipse Ti-E inverted microscope fitted with 23, 103, and 203 objectives. For laser scanning confocal microscopy, anesthetized larvae were embedded in 1.5% low-melting-point agarose on optical bottom plates or dishes (MatTek Corporation). For long term microscopy, agarose was covered with fish water containing 0.007% Tricaine. A Nikon A1 confocal microscope with a 203 Plan Apo 0.75 NA objective was used to generate 35-40 μm z stacks consisting of 0.3-2 μm optical sections. The galvano scanner was used for all static imaging and for time-lapse imaging of the CHT. Time-lapse images were taken at different intervals for 5 hr. Data were acquired with NIS Elements (Nikon).

#### Embryo survival assay

In vitro-transcribed mRNA’s for venus, venus-2A-baxa, venus-2A-ΔBH3-baxa and venus-2AMOM-baxa were diluted in injecting solution and injected into the yolk of one- to two-cell-stage embryos (Tobin et al., 2012) at a concentration of 300ng/μl. Embryo survival was monitored 5 and 24 hours post injection.

#### Assessment of camptothecin-induced apoptosis in whole zebrafish larvae

2 dpf larvae were treated by bath with 500 nM camptothecin or DMSO (0.5% final concentration) and incubated for 6 hours at 28.5°C (Langheinrich et al., 2002). Then all larvae from each treatment were transferred to 1.5 ml centrifuge tubes containing 1 ml of fish water and 2.4 μg/ml acridine orange (Sigma) and incubated protected from the light with rotation at room temperature for 30 minutes (Paquet et al., 2009). Finally, larvae were washed twice in fish water with tricaine and acridine orange fluorescence was analyzed by microscopy as readout for apoptosis. ImageJ was used to quantify mean grey value for each animal as indicated in supplemental figure 1.

#### Mitochondrial Ca2+ overload detection and quantification assay

Mitochondrial Ca2+ overload was assayed by the increase in fluorescence intensity of the cell permeant mitochondrion-targeted Ca2+ indicator Rhod-2 AM (Pozzan and Rudolf, 2009) (Fisher Scientific) or the genetic encoded Ca2+ reporter GCaMP3 targeted to the mitochondrion [Tg(BH:GFP-mfap4:Mito-GCaMP3)]. Tg(mpeg1:YFP)^w200^ or Tg(mpeg1:Brainbow)^w201^; Tg(BH:GFP-mfap4:Mito-GCaMP3) larvae were infected with 90-120 EBFP2-expressing Mm. To calculate the percentage of macrophages positive for Rhod-2, 1 dpi Tg(mpeg1:YFP)^w200^ animals were microinjected via CV with a solution containing TNF and 31.25 μg/ml Rhod-2 or a solution containing vehicle for TNF and Rhod-2 only. 50 μg Rhod-2 were diluted in 20 μl DMSO and stored in 1 μl aliquots at −20°C and protected from light. Then Rhod-2 was diluted 1:40 in PBS and added 1:2 to the TNF solution. To quantify increased mitochondrial Ca2+ concentration, 1 dpi Tg(mpeg1:Brainbow)^w201^; Tg(BH:GFP-mfap4:Mito-GCaMP3) animals were microinjected via CV with TNF. After TNF or TNF and Rhod-2 administration, larvae were prepared for time-lapse confocal imaging. Time-lapse images were taken at different intervals for 5 hr. Mitochondrial GCaMP3 fluorescence was quantified as maximum fluorescence intensity per macrophage using NIS-Elements.

#### Protein sequence analysis

LALIGN (Myers and Miller, CABIOS 1989) (https://embnet.vital-it.ch/software/LALIGN_form.html) was used for global analysis without end-gap penalty of aminoacid residue sequences. Percentage identity, percentage similarity and global/local score are shown in Table 1. Protein accession numbers: hBAX alpha (NM_138761.3), hBAX beta (NM_004324.3), hBAK1-202 (NM_001188.3), hBAK1-203 (ENST00000442998.6), zfBAXA (NM_131562), zfBAXB (NM_001013296).

#### Mitochondrial ROS quantification assay

Mitochondrial ROS production was assayed by fluorescence intensity of the cell permeant mitochondrion-targeted MitoTracker Red CM-H_2_-Xros (Roca and Ramakrishnan, 2013) (Fisher Scientific). Tg(mpeg1:YFP)^w200^ larvae were infected with 90-120 EBFP2-expressing Mm. Animals were microinjected 1 dpi via CV with a solution containing TNF and 50 μM MitoTracker Red CM-H_2_-Xros. After TNF or TNF and MitoTracker Red CM-H_2_-Xros administration, larvae were prepared for confocal imaging. A single time point image was taken at 40 minutes after TNF administration. MitoTracker Red CM-H_2_-Xros fluorescence was quantified as maximum fluorescence intensity per macrophage using NIS-Elements.

### QUANTIFICATION AND STATISTICAL ANALYSIS

The following statistical analyses were performed using Prism 5.01 (GraphPad): One-way ANOVA with Bonferroni’s post-test, Fisher’s exact test and Student’s unpaired t test. Error bars represent standard error of mean. Post-test *P* values are as follows: Not significant, * p < 0.05; ** p < 0.01; *** p < 0.001. The statistical tests used for each figure can be found in the corresponding figure legend. Where the *n* value is given and not represented graphically in the figure, *n* represents the number of zebrafish used for each experimental group.

### DATA AND SOFTWARE AVAILABILITY

The following software was used: NIS-Elements (image acquisition in wide-field and confocal microscopy), Corel Draw (figure preparation) and ImageJ (quantification of camptothecin-induced apoptosis in zebrafish larvae); see Key Resources Table for more information.

## TABLE FOR AUTHOR TO COMPLETE

Please upload the completed table as a separate document. **<u>Please do not add subheadings to the Key Resources Table.</u>** If you wish to make an entry that does not fall into one of the subheadings below, please contact your handling editor. (**NOTE:** For authors publishing in Current Biology, please note that references within the KRT should be in numbered style, rather than Harvard.)

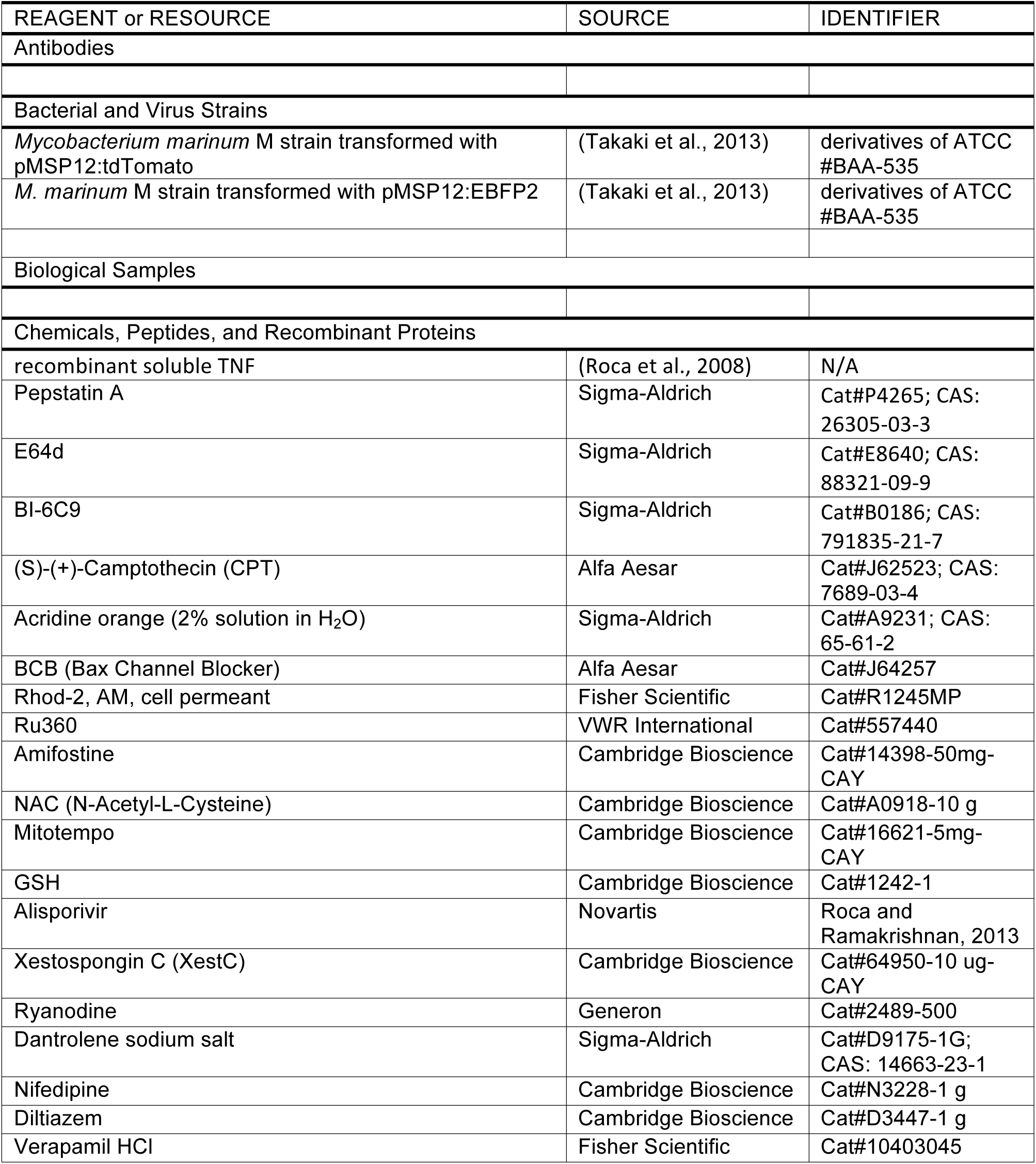

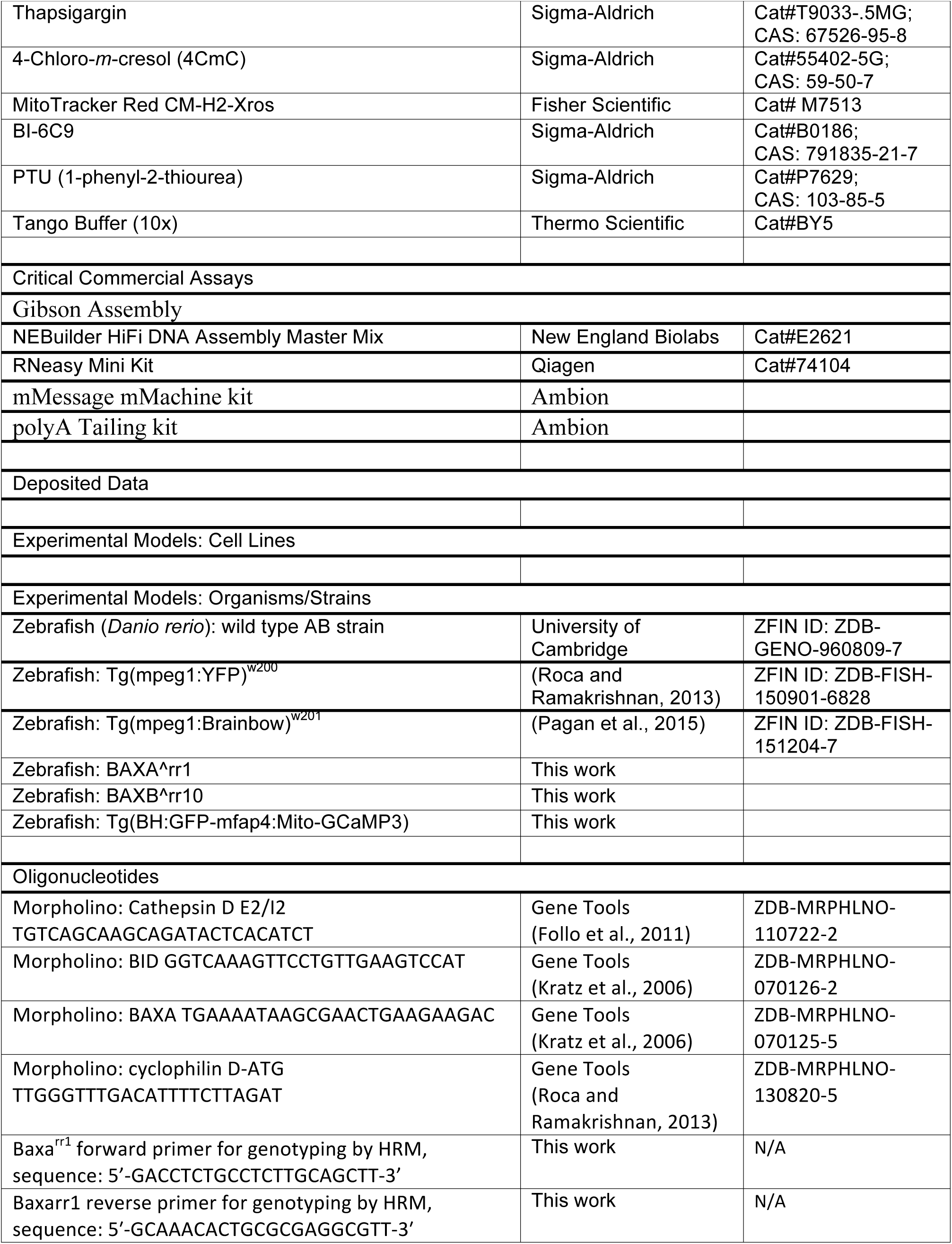

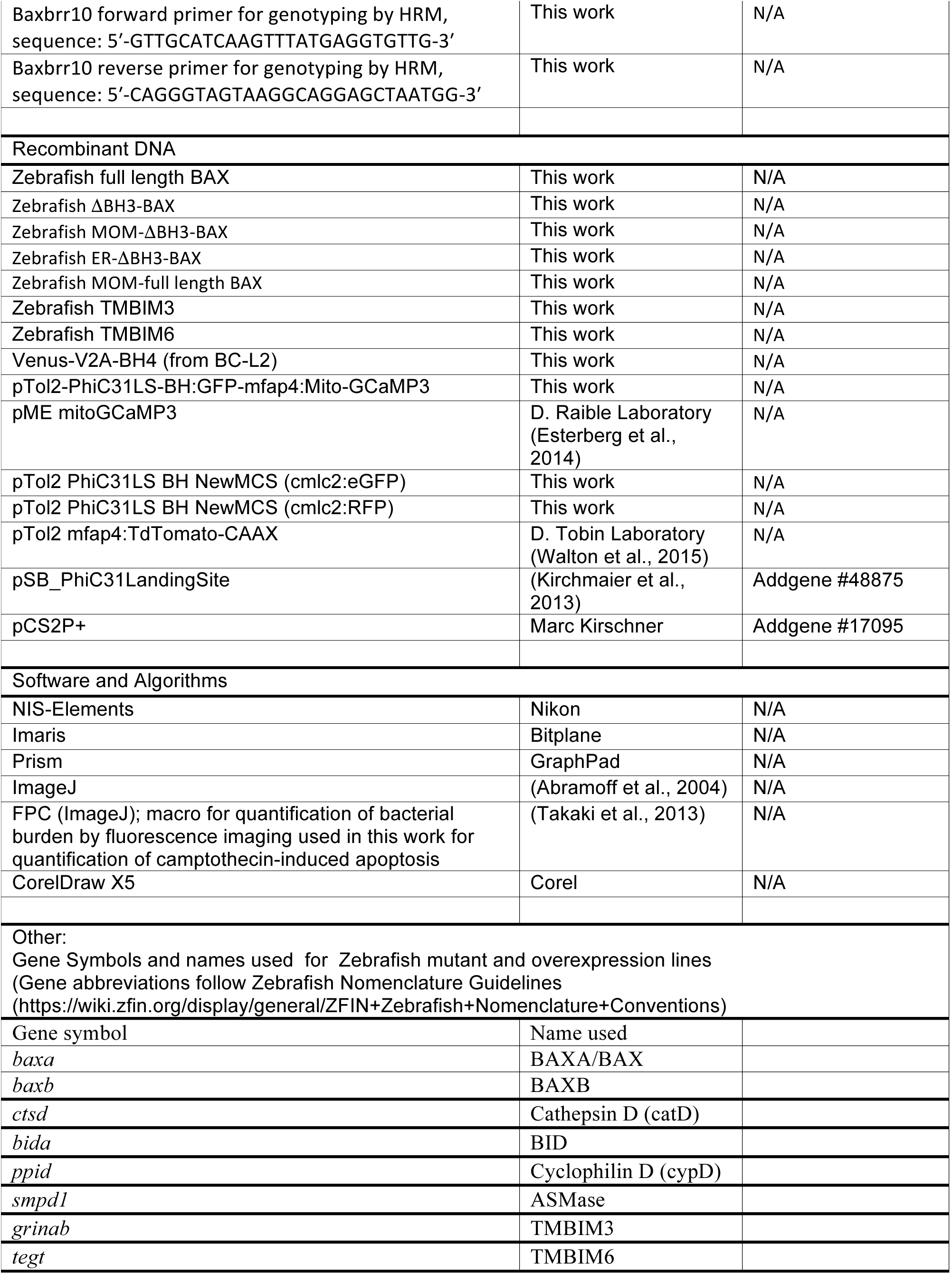

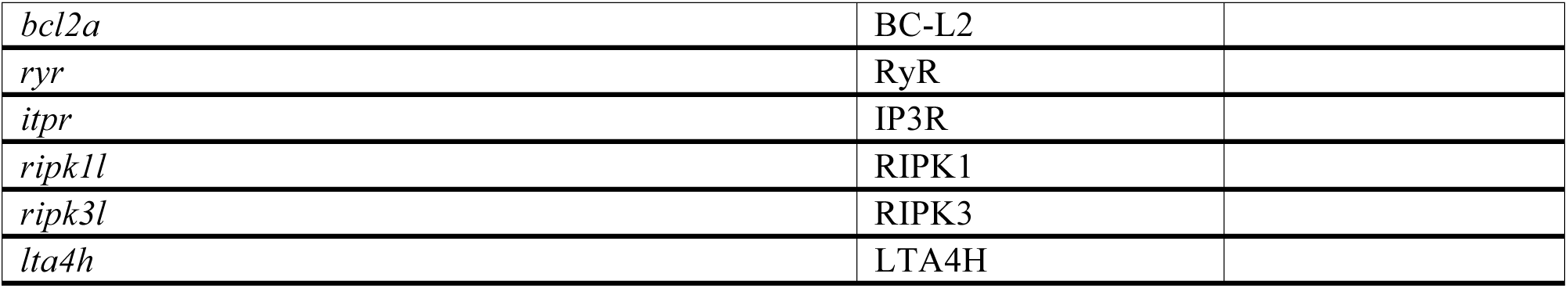
KEY RESOURCES TABLE.

